# A unified central thalamus mechanism underlying diverse recoveries in disorders of consciousness

**DOI:** 10.1101/2024.11.04.621844

**Authors:** Haoran Zhang, Qianqian Ge, Xiao Liu, Yuanyuan Dang, Long Xu, Yutong Zhuang, Si Wu, Steven Laureys, Jianghong He, Shan Yu

## Abstract

Disorders of consciousness (DoC) encompass a range of states characterized by prolonged altered awareness due to heterogeneous brain damage and are associated with highly diverse prognoses. However, the neural mechanisms underlying such diverse recoveries in DoC remain unclear. To address this issue, we analysed neuronal spiking activities recorded from the central thalamus (CT), a key hub in arousal regulation, in a cohort of 23 DoC patients receiving deep brain stimulation treatment. Using machine learning techniques, we identified a core set of electrophysiological features of the CT, particularly the theta rhythm, that could account for individual recovery outcomes across highly varied etiologies (trauma, brainstem hemorrhage, and anoxia), clinical baselines and patient ages. These features also correctly identified one subgroup of patients who exhibited poor initial clinical manifestations but recovered unexpectedly. Simulating a conductance-based model further revealed the neurodynamics of the theta rhythm in the CT during different stages of consciousness recovery. Taken together, these findings uncover a previously unknown, unified CT mechanism that governs the recoveries in DoC.

## Introduction

Disorders of consciousness (DoC) encompass conditions characterized by impaired arousal and awareness due to severe brain damage, including coma^1^, unresponsive wakefulness syndrome/vegetative state (UWS/VS)^2–4^, and minimally conscious state (MCS)^5,6^. These conditions impose significant challenges to patients’ families and the healthcare system due to the difficulty of managing prolonged bedridden status, demanding care, and low rates of functional recovery^3,7^. Importantly, the causes and extent of brain damage leading to DoC are highly heterogeneous, including trauma, stroke, and anoxia, and ranging from focal injuries such as brainstem hemorrhage (BSH) to extensive lesions across the cortex^8^. Heterogeneous etiologies are associated with varying prognoses. For instance, individuals with traumatic injuries often exhibit better outcomes compared to those with anoxic brain injuries^4,7,9–12^. Factors such as the clinical baseline (e.g., UWS, MCS), age, and time since damage onset further contribute to variations in prognoses^3,11–14^. However, the neural mechanism underlying such diverse recoveries in various clinical factors remains unclear.

Non-invasive studies using fMRI and EEG have suggested that the status of macroscopic brain networks, including the default mode network, executive control network, and thalamocortical connectivity integrity^15–17^, as well as neuronal oscillation and synchrony within the fronto-parietal region^18–20^, may correlate with DoC recovery. However, the role of the thalamus, particularly the central thalamus (CT), in functional recovery in DoC remains incompletely understood as its activities cannot be assessed with high resolution using non-invasive methods. As a bidirectional hub in arousal regulation, the CT receives inputs from the brainstem through the ascending reticular activating system (ARAS)^21–23^ and sends broad projections modulating the frontal-striatal system^13,21,22,24–29^. Despite its critical role, direct recordings from the CT in DoC patients are scarce, with studies limited to a few cases or only coarse group-level statistical comparisons^30–32^.

In this study, we recorded CT activities in 23 DoC patients with heterogeneous etiologies: trauma (N=6), BSH (N=8), and anoxia (N=9), all of whom had relatively well-preserved brain structures (TableS1 and FigureS1). Among them, eight patients demonstrated consciousness recovery (CR) after one year of CT-DBS treatment, characterized by the regaining of language processing capabilities. We investigated two modalities of CT activity—neuronal spiking and multiunit activity (MUA)—to identify key electrophysiological features of CT that could predict individual recovery outcomes. These findings suggest that the CT’s activities, particularly those related to theta band oscillations, play a crucial role in functional recovery across various clinical factors in DoC.

## Results

### CT features enable individualized prognosis for DoC

All patients underwent deep brain stimulation (DBS) treatment targeting the CT (Figure1A). We investigated two modalities of CT activity, including neuronal spiking and multiunit activity (MUA) (Figure1B). Compared to the local field potential, which serves as a proxy for synaptic input and is significantly confounded by volume conduction, MUA provides more localized information regarding population outputs within a range of approximately 200–300 μm from the recording electrode^33,34^, which has demonstrated unique strength in characterizing activities of small nuclei^35,36^. Leveraging these two modalities, we examined a total of 34 electrophysiological features to characterize the CT state, covering multifaceted properties including the discharge properties of single neurons^26,37^, synchronization among neuron populations^35,36,38^, coherence between the activity of individual neurons and that of population^26,27,39,40^, and background noise levels^41^.

**Figure1.**
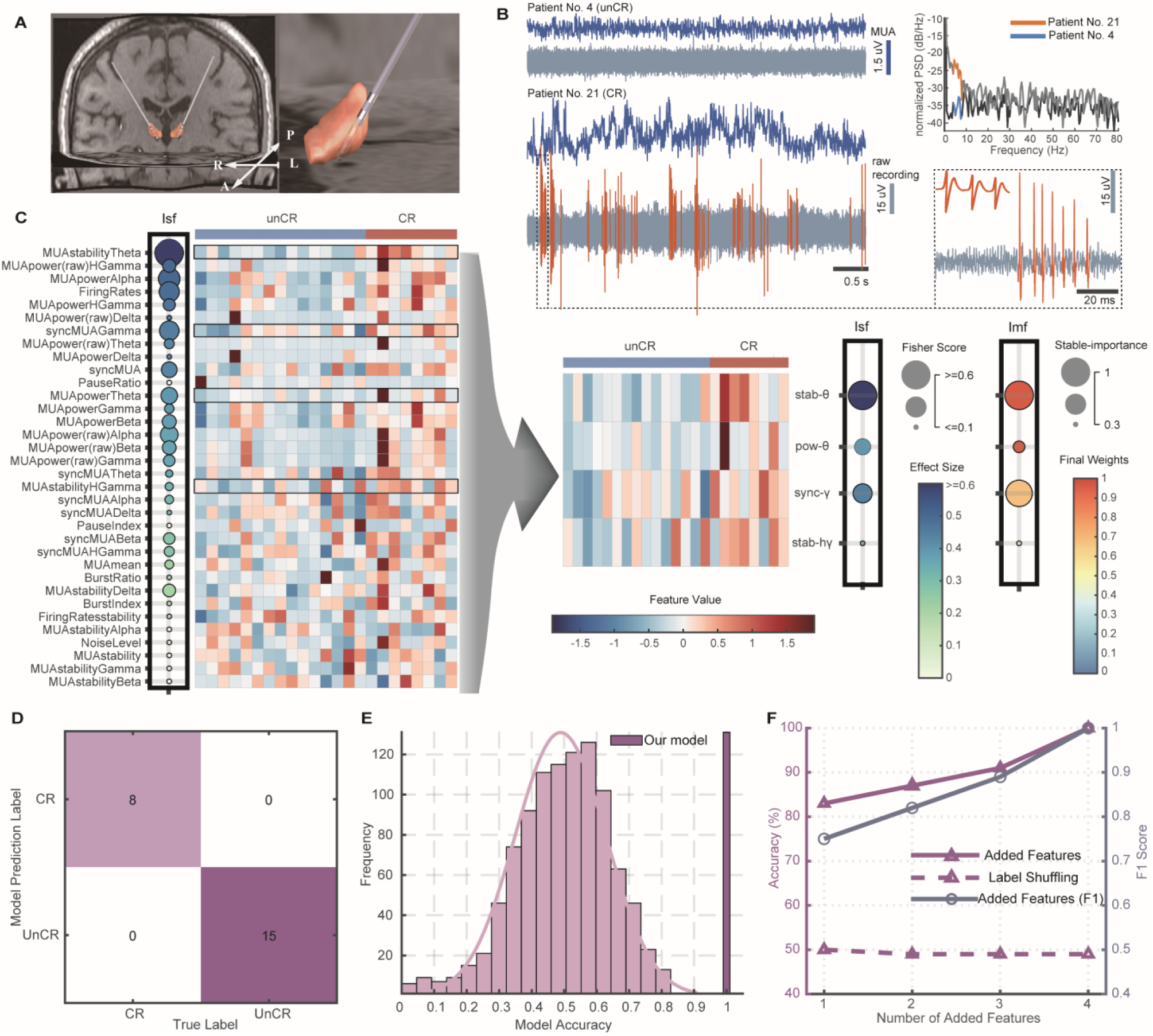
Individualized prognosis for DoC using CT features. **(A)** MRI coronal view showing electrode placements in the bilateral central thalamus of an example patient (Patient No. 18), with an inset showing the left hemisphere electrode. **(B)** Visualization of electrophysiological signals extracted from the raw recording (grey): neuronal spikes (orange) and multi-unit activity (MUA, blue). Examples from two patients are shown: Patient No. 4 (unCR) is characterized by a predominantly silent state, exhibiting only background activity, while Patient No. 21 (CR) displays intermittent bursting activity and corresponding unstable MUA activity. The normalized power spectrum (PSD) for the MUA signals of both patients is presented, with the theta band highlighted. **(C)** Electrophysiological profiles of the CT before and after feature selection: The left panel shows the distribution of 34 CT features across patients. The importance of Single-Feature (Isf) is indicated by the Fisher Score (circle size) and effect size (color intensity), illustrating the variability among individual features. The four selected features are highlighted with black borders and are further detailed in the right panel, including stab-θ, pow-θ, sync-γ, stab-hγ. The importance in the Multi-Feature Combination (Imf) represents the stable-importance index (circle size) and final feature weights (color intensity) in the CT metric (FigureS4A). These four features are visualized using representative data from two groups (blues/red for unCR/CR). Pow-θ is represented by the amplitude of the theta band oscillation; stab-θ and stab-hγ are visualized through the stability of the shown envelopes of theta and high-gamma band oscillations, respectively, and sync-γ is depicted by the synchronization strength between gamma band MUA fluctuations (dark) and neuronal spiking (light). **(D)** Confusion matrix demonstrating accurate classification of all 23 patients (15 unCR, 8 CR). **(E)** Permutation Test: The accuracy of the model (dark pink) and that using shuffled data as the control (light pink). **(F)** Ablation study: progressive improvement in model accuracy and F1 score with more features are added (with the sequence of stab-θ, pow-θ, sync-γ, stab-hγ), contrasted with baseline performance near chance level (50%) from shuffled labeling (refer to Methods for additional details).

We found that the electrophysiological features of the CT show redundancy, noise, and variable discrimination power (Figure1C, left). To select the key features and examine their predictive power for individual recoveries^16,20,42,43^, we employed a machine learning pipeline to identify the optimal feature combination (FigureS2A). As a result, we identified a combination of four features (Figure1C, right) that could correctly discriminate between patients who exhibited CR and those who did not (unCR) (Figures1D, S3B-S3D and S4B) —a discrimination that unselected features failed to achieve (FigureS3A). These four features include: 1) MUA stability in the theta band (stab-θ), which is associated with thalamic intermittent bursting patterns (see Figure1B for an example)^44,45^ ; 2) MUA power in the theta band (pow-θ), measuring the amplitude of theta band oscillation of MUA; 3) syncMUA in the gamma band (sync-γ), measuring the synchronization of neuronal spiking and MUA population fluctuation in the gamma band; and 4) MUA stability in the high gamma band (stab-hγ). The Permutation Test showed that the accuracy of the model was significantly higher than the chance level (Figure1E), and the ablation study confirmed that all four features contributed to the model’s performance (Figure1F and TableS4).

### CT’s status provides a unified indicator of DoC recovery across diverse clinical factors

After identifying key features characterizing the CT state, we averaged the weight parameters of the selected set to develop a CT-based metric for DoC prognosis (FigureS4A). In the present dataset, we first confirmed previous findings that diverse clinical factors, including diagnostic baselines^3,10,11,13^, ages^12,14^, and etiologies^4,7,9–12^, influence prognosis. We measured the CR rate across various groups defined by these clinical factors. A higher CR rate was observed in the MCS-group compared to the UWS group (Figure2A inset, Fisher’s Exact Test, p=0.006, Odds Ratio=16.10, Cohen’s h=1.35). Similarly, the younger group demonstrated a higher CR rate compared to the older group (Figure2B inset, Fisher’s Exact Test, p=0.006, Odds ratio=16.10, Cohen’s h=1.35). Regarding etiology, the highest CR rate was observed in the traumatic group— 66.7% in trauma, 22.2% in BSH, and 25% in anoxia. However, the differences among three etiologic groups were not statistically significant (Fisher’s Exact Test, Figure2C inset). Importantly, we found that, across all these groups defined by clinical factors, the patients whose CT metric score was above a specific threshold eventually recovered (Figures2A-2C). This indicates that the CT’s status is a key indicator of functional recovery in DoC across various clinical factors, suggesting that DoC recoveries share a common mechanism in which the CT plays a pivotal role.

**Figure2.**
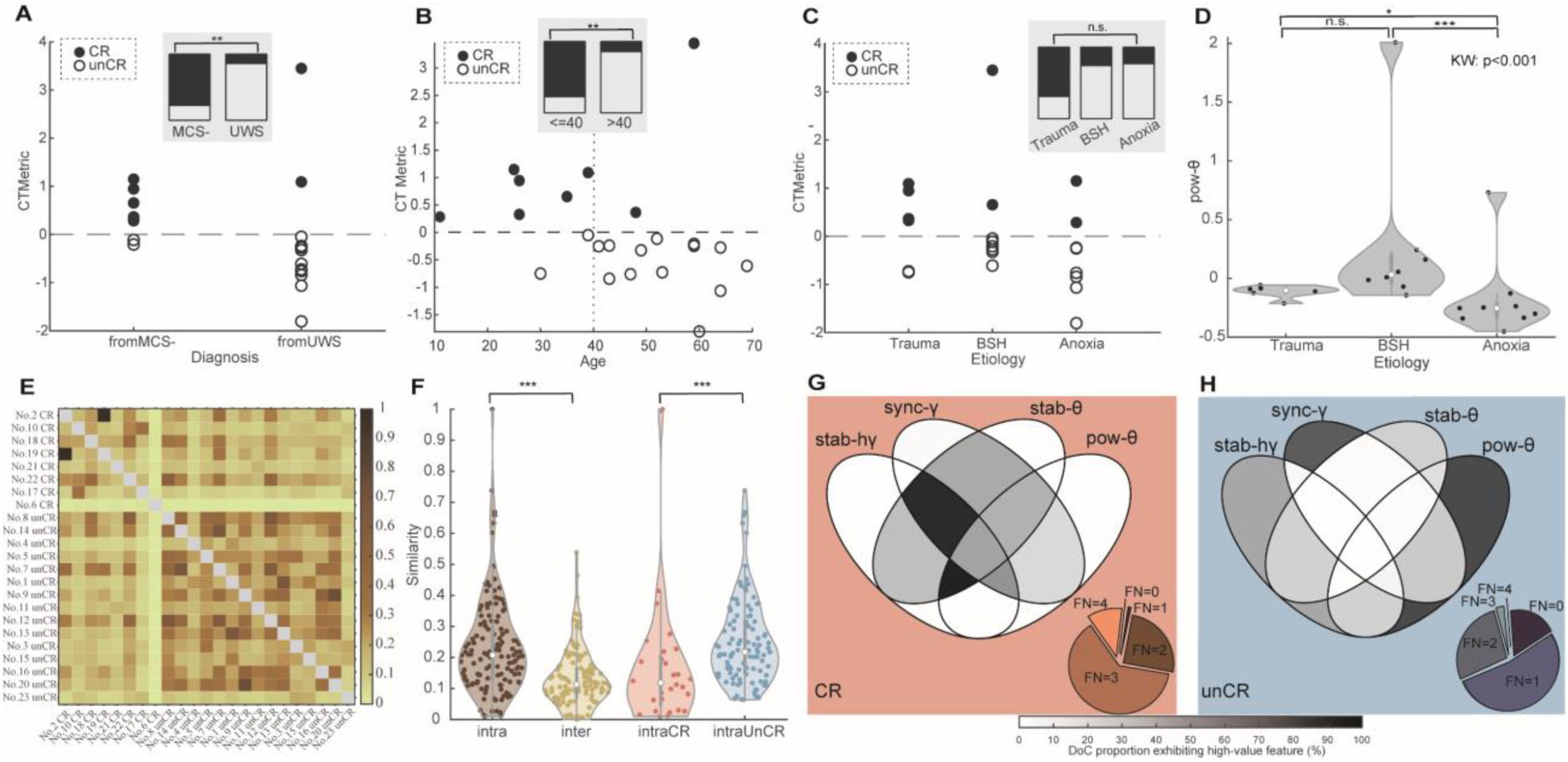
CT neurophysiological features associated with DoC prognosis (A-C), Distribution of CT metric scores across various clinical factors: (**A**) clinical baselines (MCS, UWS), (**B**) age, and (**C**) etiologies, which include trauma, brainstem hemorrhage (BSH), and anoxia. Solid circles represent CR and open circles represent unCR. The dotted line indicates the recovery threshold for the CT metric. Inset bar graphs depict the proportion of CR patients (solid) versus unCR patients (open) within each group. **(D)** The pow-θ strength of the CT among etiologies. **(E)** CT-based patient-similarity matrix: The upper left region illustrates intra-CR patient similarity, and the lower right region shows intra-unCR patient similarity. **(F)** Statistical analysis of similarities from the patient-similarity matrix, including intra-group similarity, inter-group similarity, intra-CR similarity and intra-unCR similarity. Notably, these observed statistical differences in patient-similarity are only evident with the selected CT metric and do not show significant differences prior to feature selection (FigureS5). **(G)** Venn diagram illustrates CT feature pattern in CR patients. The accompanying pie chart details the distribution of patients categorized by the number of features exhibiting high values simultaneously (feature number, FN): only 0.9% exhibit high values in a single feature; 25%, 62.8%, and 11.3% in two, three, and four features (FN=1,2,3,4), respectively, with no patients exhibiting low values in any features (FN=0). **(H)** Venn diagram and pie chart depict the CT feature pattern in unCR patients with a tendency of high values in only a single feature (FN=1), predominantly sync-γ or pow-θ. A majority of unCR patients (53.1%) show high values in just one feature, with 27.4% and 2.6% showing high values in two and three features (FN=2,3), respectively. None exhibit high values coherently across four features (FN=4), but 16.9% showing low values in all features (FN=0).

Additionally, we examined etiological differences for each selected CT feature separately. Only pow-θ showed significant discrimination among etiologies (Figure2D, Kruskal–Wallis (KW) Test, p<0.001, η2=0.51). Post-hoc tests (Conover’s All-Pairs Rank Comparison Test) revealed significantly stronger pow-θ in the BSH group compared to anoxia (p<0.001) and in the traumatic group compared to anoxia (p=0.042). Although there was a trend toward higher pow-θ in BSH compared to trauma, the difference was not significant (p=0.056).

To better understand the different CT statuses that may lead to varying prognoses, we examined the differences that discriminate between the CR and unCR groups. The patient-similarity matrix based on the CT metric (Figure2E) shows that the intra-group similarity is significantly higher than the inter-group similarity (Figure2F). In addition, intra-group similarity is significantly lower in the CR group than in the unCR group (Figure2F), indicating that unCR patients exhibit more consistent electrophysiological patterns. Additionally, we analyzed activity patterns of the CT metric in the CR and unCR groups, respectively. Venn diagrams illustrate distinct feature distributions: CR patients demonstrated higher values across multiple features simultaneously (Figure2G), while unCR patients generally exhibited lower activity levels, with typically only a single feature showing high values (Figure2H).

### CT features discriminate two subgroups of CR patients

The reduced patient similarity within the CR group suggests the presence of more nuanced classes beyond the simple dichotomy of CR and unCR. Next, we examined this possibility using unsupervised hierarchical agglomerative clustering based on CT features to search for possible subgroups. As a result, we identified three subgroups (Figures3A, S6A and S6B).

**Figure3.**
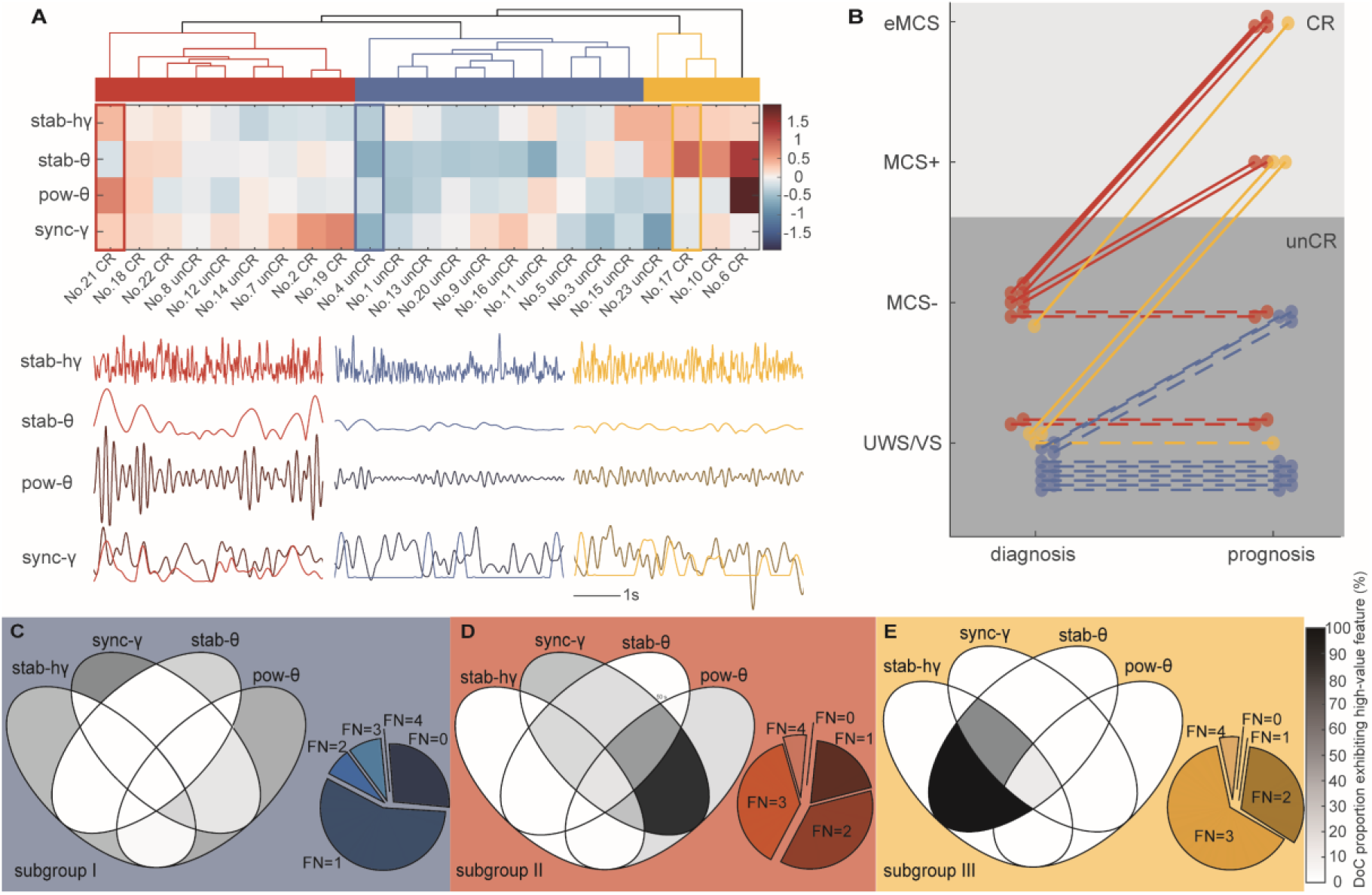
CT-based subgrouping of DoC patients and their implications for consciousness states. **(A)** Hierarchical agglomerative clustering of CT features categorizes DoC patients into three subgroups (Ⅰ in blue, Ⅱ in red, Ⅲ in yellow), each column representing a patient and each row a CT feature. Visualization of four features in an example from each subgroup: Patient No. 21 (Ⅱ), No. 4 (Ⅰ), and No. 17 (Ⅲ). Patient No. 21 (Ⅱ) exhibits stronger theta band oscillation (pow-θ) and increased synchronization between gamma band MUA fluctuations (dark red) and neuronal spiking (red), indicating enhanced sync-γ. Patient No. 17 (Ⅲ) shows both higher stab-θ and stab-hγ, illustrated by the stability of the displayed envelopes corresponding to oscillations. Conversely, Patient No. 4 (Ⅰ) exhibits weaker theta rhythms (pow-θ), reduced and even negative sync-γ, and more pronounced instability in the envelopes of oscillations (stab-θ and stab-hγ). **(B)** Changes in consciousness states pre- and post-CT-DBS therapy across 23 patients, color-coded by subgroup. Solid lines indicate patients who improved to CR after treatment, while dashed lines represent those who remained unCR. All CR patients exhibited significant improvements after CT-DBS treatment. **(C-E)** Venn diagrams and accompanying pie charts depict CT feature profiles in each subgroup. Subgroup Ⅰ mainly shows low feature values (**C**). Subgroup Ⅱ predominantly exhibits simultaneous high values in pow-θ and sync-γ (**D**), while Subgroup Ⅲ is characterized by simultaneous high values in stab-θ and stab-hγ (**E**). The pie charts detail the distribution of patients categorized by the number of features exhibiting high values concurrently (feature number, FN): Subgroup Ⅰ—56.7% with one feature, 7.2% with two, 8.6% with three, 0% with four features (FN=1,2,3,4) and 27.5% showing low values in all features (FN=0) (inset in (**C**)). Subgroup Ⅱ—19.3% with one, 36.8% with two, 37.9% with three, 6% with four, and 0% with zero (FN=1,2,3,4,0, inset in (**D**)). Subgroup Ⅲ—0% with one, 31.8% with two, 63.2% with three, 5% with four, and 0% with zero (FN=1,2,3,4,0, inset in (**E**)).

Subgroup Ⅰ exhibited overall lower values across all four features (Figures3C, S6C and S6D). Subgroup Ⅱ featured higher pow-θ and sync-γ (Figures3D, S6C and S6D). Subgroup Ⅲ was characterized by higher stab-θ and stab-hγ (Figures3E, S6C and S6D). We found that none of the patients in Subgroup Ⅰ exhibited recovery. That is, all CR patients were exclusively in subgroups Ⅱ and Ⅲ. Statistical tests confirmed that Subgroups Ⅱ and Ⅲ exhibited a higher CR ratio than Subgroup Ⅰ (Fisher’s Exact Test, p=0.005 for three-group differences; Subgroup Ⅰ vs. Ⅱ, p=0.011, Odds ratio=0, Cohen’s h=-1.68; Subgroup Ⅰ vs. Ⅲ, p=0.011, Odds ratio=0, Cohen’s h=-2.09). Taken together, these results suggest that the CR group could be further divided into two classes, each with distinct CT features.

In addition, these three subgroups showed significant differences in their initial clinical baselines regarding UWS or MCS-(Fisher’s Exact Test, p=0.018). Subgroup Ⅰ consisted exclusively of UWS patients. Subgroup Ⅱ primarily consisted of MCS-patients (7 out of 9). Subgroup Ⅲ predominantly comprised UWS patients (3 out of 4) (Figures3B and S6E). Interestingly, in this cohort, only two patients recovered consciousness from the UWS state, both of whom belonged to Subgroup Ⅲ (TableS1). Statistical analyses revealed no significant difference in age or etiology among three subgroups. In summary, CT features differentiate two subgroups of patients who exhibit favorable recovery potential, with notable differences in their recovery modes. Subgroup Ⅱ patients recovered exclusively from the MCS-state, whereas Subgroup Ⅲ patients showed surprising recovery from the UWS state.

### Simulation of neural dynamics underlying theta rhythms in the CT of DoC patients

The features of theta rhythms in the CT, especially stab-θ and pow-θ, were identified as the most significant contributors to our prediction model (TableS4, see also Eq. 9 in Methods). These theta/alpha rhythms, indicative of the low-threshold bursting (LTB) regime mediated by T-type calcium channels, are well-established neurophysiological characteristics^46–49^ in the thalamus during altered consciousness under anesthesia and sleep^44–46,50^. Our finding of the significance of theta rhythms suggested a similar LTB regime in DoC. To examine this issue, we developed a biophysical model for CT activity that incorporates DoC-specific physiological conditions and investigated whether it can generate theta rhythms. The model included three thalamic nuclei— thalamic reticular nucleus (TRN), CT, and specific relay nuclei (SR) (Figures4A and 4B)— crucial for modulating awareness/arousal^21,22,25,45,50–53^. Under normal conditions of both membrane potential and afferent inputs^54^ (Figure4C), the model exhibited desynchronized tonic firing activity, similar to that observed during healthy wakefulness in the thalamus^8,55,56^ (Figure4D).

**Figure4.**
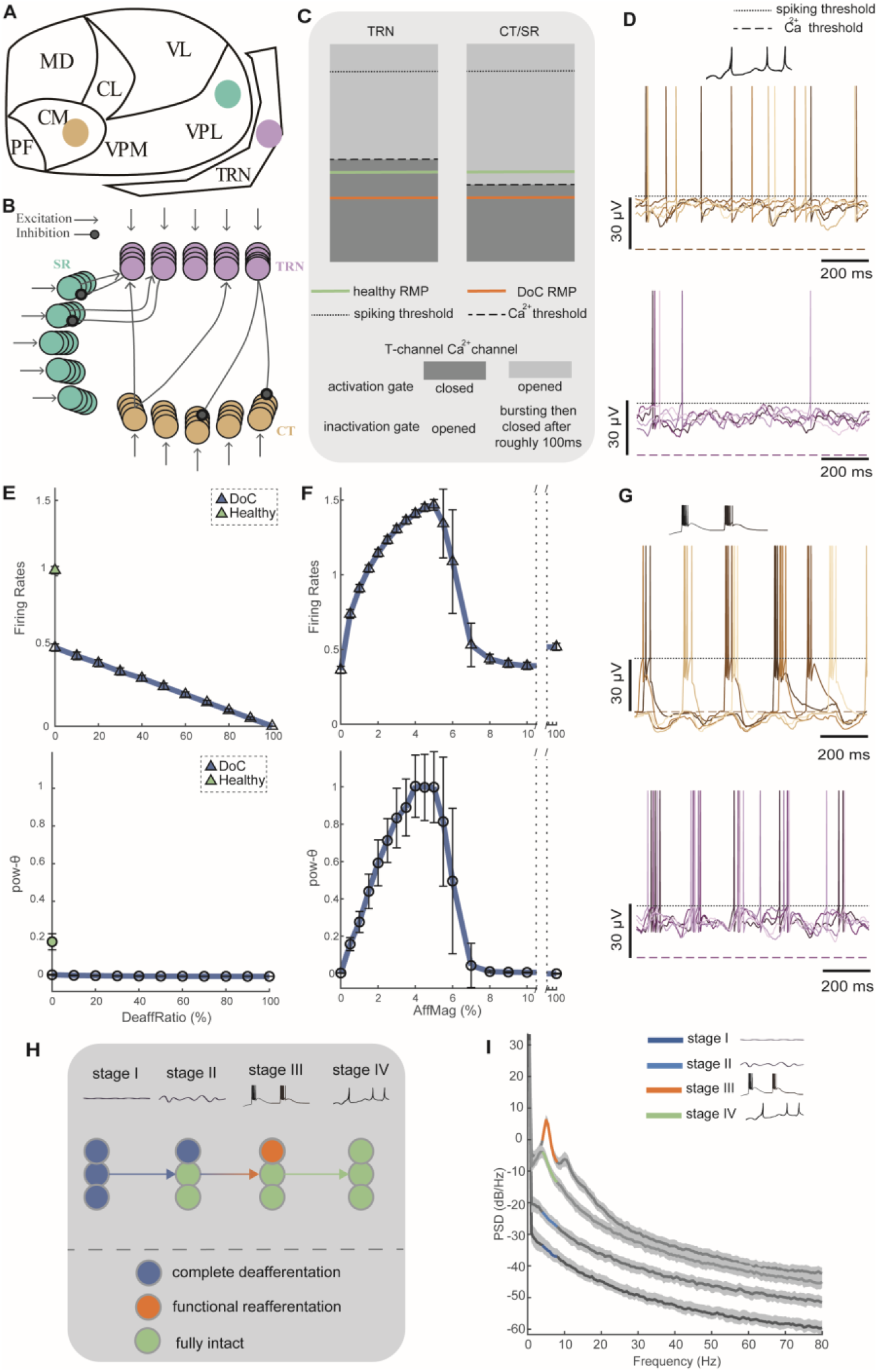
Network simulation of neural dynamics underlying theta rhythms in CT. **(A)** Schematic of thalamic nuclei (coronal view): CT (amber), thalamic reticular nucleus (TRN), and specific relay nuclei (SR). The SR encompasses first-order sensory relay nuclei such as the ventral posterior nucleus (somatosensory), the medial geniculate nucleus (auditory), and the lateral geniculate nucleus (visual). **(B)** Network connectivity diagram illustrating that the SR is wired to the TRN with one-to-one connections, and the CT is linked to the TRN with random one-to-many connections (see TableS5 for details). **(C)** Diagram of thalamic neuronal parameters, including the healthy and DoC resting membrane potential (RMP, solid line), spiking threshold (dotted line), and the threshold for T-type calcium channels (dashed line, see TableS6 for details). Specifically, when the membrane potential is initially below the calcium threshold, the calcium inactivation gate remains open. Once the potential exceeds this threshold, the calcium activation gate also opens, leading to a low-threshold bursting regime for roughly 100 ms before the inactivation gate closes. **(D)** Tonic regime traces from five example CT neurons and five TRN neurons, with raster plots in the FigureS7A. **(E)** State transitions under complete deafferentation (no residual input for affected neurons) are demonstrated by varying proportions of deafferented neurons (deaffRatio), and measured through changes in firing rates and theta band power (pow-θ). The healthy profile (green) is also depicted, characterized by a deaffRatio of 0% and the restoration of normal RMP. **(F)** State transitions under reafferentiation with various magnitudes of residual afferent inputs (affMag). The firing rate and pow-θ are shown during transition from complete deafferentation to a theta rhythmic state, and then to further reafferentiation. The deaffRatio is set at 30%, a level that would not significantly affect rhythmic properties. **(G)** Bursting regime traces from five CT neurons and five TRN neurons, with raster plots in FigureS7B. **(H)** Diagram illustrating the CT status across four consciousness stages (see Methods for details). **(I)** Power spectral density (PSD) of CT across four consciousness stages, depicted by color transitions from dark blue (Stage Ⅰ) to blue (Stage Ⅱ), orange (Stage Ⅲ), and green (Stage IV) for theta band power, with standard deviations shown in light gray.

The pathophysiological mechanism underlying DoC involves the reduction of excitatory synaptic activity, attributed to structural/functional deafferentation and a lowered neuronal resting membrane potential (RMP)^8,44,55,56^. Thus, we simulated CT activity under lower RMPs (Figure4C) and various afferent conditions. We found that merely increasing the proportion of deafferented neurons (deaffRatio) induced a quiescent state with marginally reduced firing rates, but did not introduce theta rhythms in the CT (Figure4E). However, additionally augmenting the magnitude of residual afferent input (affMag) in those affected neurons shifted neuronal activity toward an LTB regime (Figure4G), facilitating the emergence of theta rhythms, accompanied by a similar change in firing rates (Figure4F). These results reveal how the theta rhythmic state is generated in the CT of DoC patients.

To further simulate the CT state in different stages of DoC recovery, we varied RMPs, deaffRatio, and affMag in the model. Stage Ⅰ features deafferentation in more than 90% of neurons in the CT; Stage Ⅱ involves deafferentation in 30% of CT neurons; Stage Ⅲ represents a theta rhythmic intermediate state; and Stage IV marks full recovery to a healthy profile (Figure4H, see Methods for details). During the transition from Stage Ⅲ to IV, our simulations revealed that both full reafferentiation and restoration of membrane potential are necessary for the CT to finally return to a tonic regime (Figures4E and 4F). Conversely, for the transition from Stage Ⅱ to Ⅰ, the decline in neurological condition resulted from increasing deaffRatio and led to a further reduction in CT firing rates (Figure4E). Collectively, these findings across four stages are consistent with the distinct CT dynamics as outlined in the ‘ABCD’ model^55,56^, which is a theoretical framework delineating CT activity patterns under varying levels of consciousness. Additionally, these findings reveal a biphasic pattern of theta rhythms, characterized by an initial emergence (Ⅱ to Ⅲ) followed by a decline (Ⅲ to IV) during functional recovery (Figure4F). This biphasic pattern, also evident in the power spectral density of the simulated CT activity across these four stages (Figure4I), mirrors a similar, biphasic pattern of theta/alpha rhythm during anesthesia^44,57^, indicating a potential shared mechanism underlying different levels of consciousness during loss of consciousness (LOC) under DoC and anesthesia.

### Neural dynamics underlying different recovery modes of DoC

As demonstrated above, two subgroups within our cohort, both with favorable recovery potential, displayed distinct CT theta rhythm profiles, characterized by higher pow-θ (Ⅱ) and stab-θ (Ⅲ), respectively (Figures3D and 3E). This variation suggested that pow-θ and stab-θ are largely independent parameters of the theta rhythm for measuring different CT states in DoC. Additionally, studies have shown that neuronal activities become increasingly unstable in the thalamocortical circuit as the level of consciousness diminishes during LOC under deep anesthesia. These unstable activities were driven by slow oscillations and/or burst suppression, either of which significantly altered the membrane potential^44,45,58,59^. Thus, to closely examine the effects of both power and stability of the theta rhythm, we simulated the CT state transition from the quiescent state (Stage Ⅱ) to the theta rhythmic state (Stage Ⅲ). Specifically, we varied both affMag and the stability of afferent inputs (affStab) to CT neurons (Figure5A).

**Figure5.**
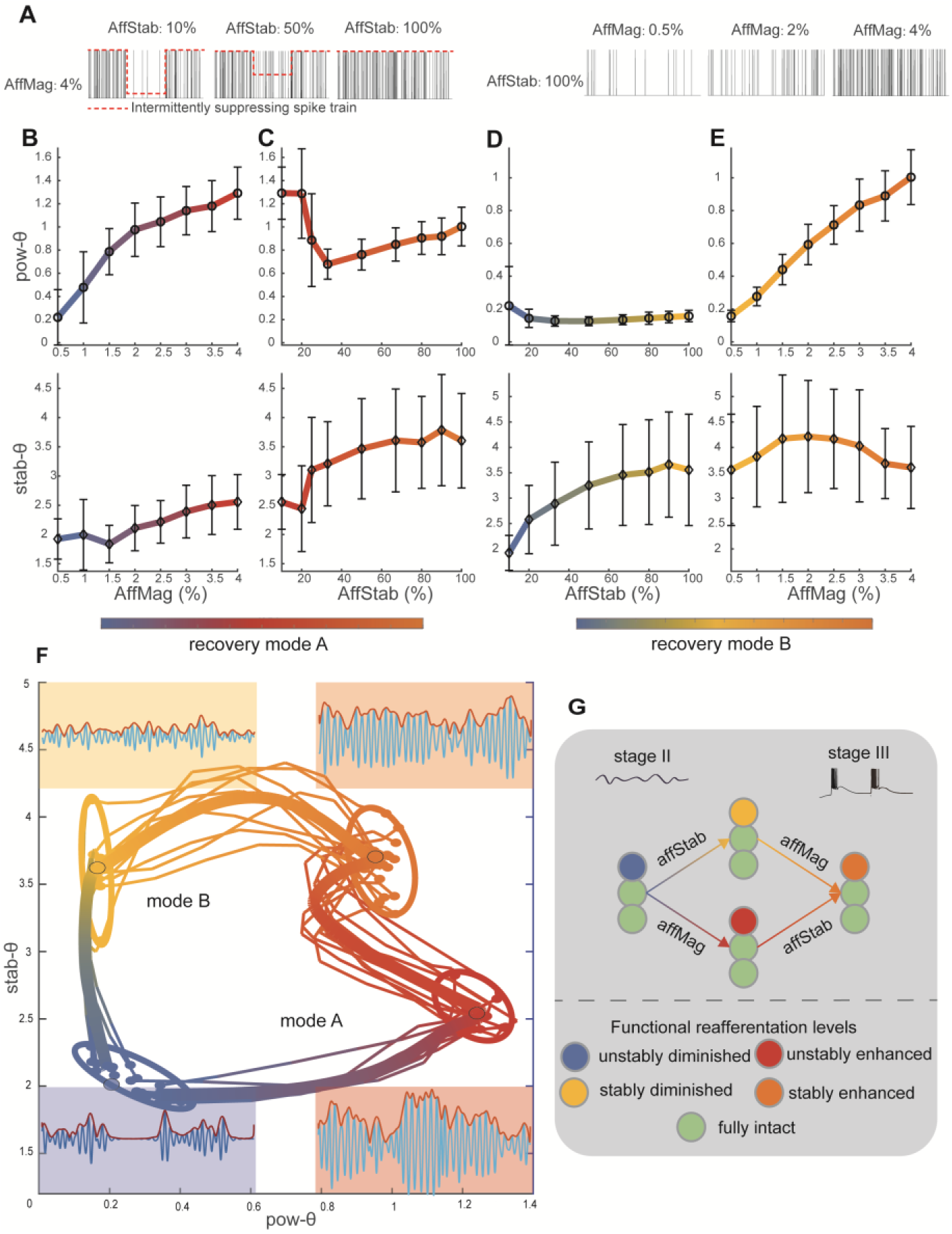
Two modes of neural dynamics underlying DoC recovery. **(A)** Illustration of residual input simulation. Left: Input stability manipulation through “OFF periods,” with afferent stability (affStab) ranging from 10% (90% of spikes muted during OFF periods) to 100% (no suppression) while afferent magnitude (affMag) remains constant. Right: Three examples of Poisson spike trains with varying discharge probabilities. AffMag ranges from 0.5% to 4%, with affStab consistently at 100%. **(B-E)** State transitions from a quiescent state (Stage Ⅱ) to a rhythmic state (Stage Ⅲ), illustrating the effects of varied afferent conditions on CT neurons. ‘Varied afferent conditions’ encompass variations in both magnitude (affMag) and stability (affStab) of residual inputs to CT. Panels (**B**) and (**E**) assess transitions under varied affMag, while panels (**C**) and (**D**) focus on transitions under varied affStab. For each subplot, the top panels display pow-θ, and the bottom panels show stab-θ. Recovery Modes A and B are indicated by color transitions from blue to red to orange (**B,C**) and blue to yellow to orange (**D,E**), respectively. Error bars represent standard deviations. **(F)** Depiction of two recovery neurodynamic modes in the feature space of the theta rhythm. Fine lines represent average trajectories of ten trials, and a thick line indicates the overall average across 100 trials. Four circled areas denote distinct states of CT: low stab-θ/pow-θ (blue), low stab-θ and high pow-θ (red), high stab-θ and low pow-θ (yellow), and high stab-θ/pow-θ (orange). Insets show theta rhythm profiles from four simulated CT examples at each corner. **(G)** Diagram illustrating the neurodynamic recovery modes as CT transitions from Stage Ⅱ to Stage Ⅲ. Five afferent conditions affecting the CT are considered.

We found that increasing affMag monotonously enhanced pow-θ while minimally affecting stab-θ (Figures5B and 5C). Similarly, affStab increased stab-θ without significantly affecting pow-θ (Figures5D and 5E). As a result, increases in affMag and affStab respectively enhanced the power and the stability of theta band activities. The changes in firing rates were in line with those of pow-θ during these processes (FigureS8). Furthermore, two neurodynamic modes could be identified (Figures5F and 5G). Mode A involves increasing the affMag first, followed by affStab, leading to an enhancement in the pow-θ first, followed by stab-θ. Conversely, Mode B follows the opposite sequence, increasing stab-θ first and subsequently pow-θ. These two neurodynamic recovery modes respectively correspond to the empirical data observed in Subgroups Ⅱ and Ⅲ (FigureS9), suggesting that they may underpin the distinct CT profiles discovered in these two recoverable subgroups (FigureS10).

## Discussion

The CT’s well-established role as a bi-directional hub in arousal regulation^21–29^ underscores its importance in exploring neural mechanisms underlying DoC. Our results confirm that CT status consistently serves as an indicator of functional recovery in patients with prolonged altered awareness across diverse clinical factors, despite considerable heterogeneity, including highly varied etiologies, clinical baselines, and ages. Notably, based on CT activities, we identified two distinct subgroups (Ⅱ and Ⅲ) of patients with different clinical baselines yet similarly favorable outcomes. Biophysical modeling demonstrated two neurodynamic pathways underlying the CT’s theta rhythmic state, each corresponding to a subgroup we observed empirically.

We found that etiological differences are primarily indicated by the pow-θ in the CT. This can be understood in light of our simulation results, which show that pow-θ is affected by the damage to the CT neurons *per se* (simulated as complete deafferentation in the model) and by the magnitude of neuronal afferent input to the CT. Anoxic patients primarily experience thalamo-cortical gray matter damage^60^. According to our simulations, damage to CT neurons *per se* led to diminished theta rhythms. Indeed, this group of patients exhibited the lowest pow-θ in our empirical results. In contrast, traumatic patients mainly experience thalamo-cortical white matter damage with diffuse axonal injury, leading to less degrees of deafferentation to CT^61^. Compared to complete deafferentation, these patients may preserve certain residual neuronal inputs and thus exhibit higher pow-θ than those in anoxia, as demonstrated by our simulations and corroborated by the patients’ data. Meanwhile, patients with BSH, retaining considerable thalamo-cortical integrity, could result in highly preserved residual afferent input to CT neurons and, consequently, the highest pow-θ among all etiologies. Taken together, these results suggest that a unified CT mechanism underlies the varying electrophysiological manifestations seen in different causes of brain damage that lead to DoC. This implies that DoC is not simply an umbrella term for separate pathologies with overlapping symptoms of impaired consciousness, despite the heterogeneous underlying causes.

Moreover, we identified two distinct subgroups (Ⅱ and Ⅲ) of patients with recovery potential. Sub<u>g</u>roup Ⅱ, characterized by high pow-θ, likely retains a high magnitude of afferent neuronal input within neurodynamic Mode A. Subgroup Ⅱ also exhibits better clinical baselines. Thus, neurodynamic mode A may underpin the pathophysiological mechanisms behind the higher recovery probabilities observed in individuals with better diagnoses^3,11^. Conversely, Subgroup Ⅲ, characterized by high stab-θ, likely preserves highly stable afferent neuronal input within neurodynamic Mode B. Despite poor clinical baselines, Subgroup Ⅲ demonstrated a favorable prognosis. Therefore, neurodynamic mode B may serve as the neurophysiological basis for another subgroup showing recovery potential, albeit with poor baselines and fewer in number^3,10,11^. Similar manifestations—poor baselines yet potential for recovery—have been previously identified in cases of covert consciousness or cognitive motor dissociation^8,62–66^. However, the relationship between our findings and these conditions remains unclear, necessitating further investigation. This divergence indicates that early clinical assessments cannot capture the neurodynamic complexity or accurately estimate the recovery potential for certain patients, which aligns with the suggestion that more information should be taken into account^66–68^.

During altered consciousness induced by sleep and anesthesia, slow oscillations accompany overall membrane potential fluctuations, resulting in alternating ON-OFF periods and interrupting stable theta/alpha rhythms^44,58,59^. As anesthesia deepens, enhanced synaptic inhibition leads neural networks to further transition towards an isoelectric state^37,69,70^. During this transition, burst suppression—a marker of deep anesthesia—intermittently mutes oscillations, leading to an increasingly unstable activity pattern^44,71^. The theta rhythm mentioned above is sustained by stable excitatory-inhibitory interactions between the central thalamus and the inhibitory TRN^45,46,49,50^. Enhanced inhibition provided by the TRN to CT cells prompts the latter to generate LTBs via T-type low-threshold calcium channels. The TRN cell, in turn, responds to this burst from CT, thus promoting a thalamic inhibitory-excitatory rhythm. However, this theta rhythm is absent when the brain remains in a quiescent isoelectric state, as observed during LOC under very deep anesthesia, resulting from a highly hyperpolarized membrane potential^69,70^. In our empirical results, the pow-θ and the stab-θ in the CT of DoC patients distinctly capture their status, ranging from quiescence to a theta rhythmic state. Additionally, dynamical simulations demonstrate that this theta rhythmic state is caused by LTB in the CT. Taken together, these results strongly suggest that the mechanism underlying LOC in DoC parallel those observed during anesthesia. By linking DoC to well-established LOC mechanisms, our findings pave the way towards a thorough understanding of DoC and its recovery processes. Given the close projection relationship between the thalamus and cortex^15,25–27,72–74^, the identification of features of theta rhythm in the CT in this study could guide the search for non-invasive prognostic rhythmic markers.

Our findings suggest that residual activities in the CT, though in only one nucleus, provide substantial information on DoC prognosis. However, it is important to recognize that all patients in this study underwent CT-DBS treatment, a promising intervention associated with higher probabilities of regaining consciousness compared to the rare instances of spontaneous recovery^10–13^. Additionally, electrical stimulation targeting the CT has shown efficacy in awakening non-human primates from anesthesia^26–28^ and treating patients with moderate-to-severe traumatic brain injury^75^. Thus, our results should be interpreted to mean that CT activities can predict recovery potential given proper treatment. Moreover, it is essential to recognize that the significance of the CT for DoC does not diminish the importance of other cerebral regions^17–20,76–78^. Further research is needed to determine whether DoC-related information is specific to CT or indicative of broader brain network activity.

## Materials and Methods

### Participants and surgical indications

39 DoC patients received CT-DBS treatment from June 2011 to June 2021 at the Seventh Medical Center of PLA General Hospital and Beijing Tiantan Hospital. The eligibility criteria for DBS surgery included the absence of extensive skull defects, no prior skull repair or shunt pump implantation, no significant surgical contraindications, and an intact bilateral thalamus structure. A comprehensive series of assessments were performed prior to surgery to evaluate the potential of regaining consciousness. These included a thorough medical history review, confirmation of the consciousness diagnosis, MRI, functional MRI, EEG, and Mismatch Negativity. Experienced clinicians assessed CRS-R scores through repeated evaluations at least three times. All findings and the experimental nature of the DBS procedure were fully discussed with the patients’ families, who made the final decision to proceed with the surgery. The inclusion and exclusion criteria of this study aligned with our prior study^32^. Briefly, all patients were diagnosed with DoC and showed no significant improvement or deterioration in consciousness for a period longer than 4 weeks. Patients with other neurological diseases or life-threatening systemic diseases were excluded.

Ultimately, 23 DoC patients (15 UWS and 8 MCS-) met the criteria and were included in this study. Clinical information and MRI details for each patient are provided in TableS1 and FigureS1. Following the principles of the Declaration of Helsinki, written consents were obtained from patients’ legal guardians or next of kin, and the study received approval from the ethics committees of PLA Army General Hospital (protocol No: 2011-0415) and Beijing Tiantan Hospital, Capital Medical University (protocol No: 2017-361-01).

### Surgical and recording procedures

We targeted the bilateral CM-pf nuclei in the central thalamus, implanting multi-channel microelectrodes (Leadpoint, Medtronic, USA) for subsequent analysis (Figure1A). Data were extracted from the signal within a 1mm range above and below the target. All procedures were performed under general anesthesia. To avoid potential influences on neuronal firing, the administration of propofol and sevoflurane was discontinued 20 minutes before microelectrode recording. Guided by the microelectrode recording, two quadripolar electrodes (PINS L302, Beijing PINS) were precisely positioned at the target sites and verified by postoperative CT or MRI.

### DBS treatment and prognosis evaluation

After the operation, the pulse generator was activated for periodic electrical stimulation at least one week after surgery. Stimulation parameters included a frequency of 100 Hz, a pulse width of 120 us, and a voltage ranging from 1.0 to 4.0 V. Clinical follow-ups extended for 12 months, and the follow-up diagnoses were determined based on the best CRS-R score attained within this period.

In the subsequent analysis of prognosis, patients finally diagnosed as UWS and MCS-were grouped as ‘consciousness non-recovery’ (unCR), indicating a sustained absence of language processing evidence (such as command-following)^5,6^. Conversely, patients who recovered to MCS+ and emergence from MCS (eMCS), categorized as ‘consciousness recovery’ (CR), showed marked improvement after CT-DBS treatment in regaining the ability to understand language or restore communications^5,6^.

### Signal processing and feature extraction

The raw electrophysiological recordings were preprocessed to extract two distinct modalities: neuronal spikes and Multiunit Activity (MUA), as depicted in Figure1B. Multiple spikes were identified in the 24 kHz-sampling frequency traces by applying the shape template^79^. To calculated the MUA, we first estimated the amplitude of population activity, which involved truncating values exceeding the mean by more than ±2 standard deviations (SDs). The root mean square (RMS) was then calculated: firstly, squaring the values to provide an instantaneous measure of spike-band power; secondly, applying a low-pass filter using a third-order Butterworth filter at 100 Hz, then reducing the sampling rate to 500 Hz; and finally, extracting the square root to obtain the RMS value. MUA outliers were subsequently identified and excluded through a moving average RMS approach, using a 20 ms window and a threshold set at 2.58 SDs^80^.

Next, we analyzed 34 electrophysiological features of the CT (Figure1C, left), categorized into five primary properties:

#### Neuronal firing properties

Including neuronal firing rates and specific patterns of activity. The degree to which spikes are temporally aggregated was assessed through two metrics. The first metric is the spike-clustering index, which is defined as the ratio of inter-spike intervals (ISIs) less than 20 ms to those over 20 ms. The second metric is the spike-clustering ratio, which compares the total duration of clustered spikes (ISI < 20 ms) to the duration of non-clustered spikes (ISI > 20 ms). Similarly, the degree to which spiking activities are suppressed intermittently was quantified using the pause index and the pause ratio. The pause index is the ratio of ISIs exceeding 100 ms to those shorter than 100 ms, and the pause ratio measures the total duration of pauses (ISI > 100 ms) against the duration without pauses (ISI < 100 ms)^81^.

#### MUA properties

Comprising the mean value of MUA (MUAmean) and both the raw and normalized power (divided by the overall power of 0.5 - 250 Hz) across six frequency bands. The frequency band includes delta (0.5 - 4 Hz), theta (4-8 Hz), alpha (8–13 Hz), beta (13–30 Hz), gamma (30–50 Hz), and high-gamma (50–80 Hz). These features are denoted as MUApower(raw)Delta, MUApowerDelta, etc.

#### Signal stability

Covering the stability of firing rates (FiringRatesstability), overall MUA (MUAstability), and stability within each specified band (MUAstabilityDelta, etc.). The signal envelopes are determined first using the Hilbert transformation. A one-second sliding window then assesses local standard deviation trends over time. The coefficient of variation (CV) quantifies the relative standard deviation to the mean, and the inverse of the CV serves as the stability strength indicator.

#### Synchroniza2tion between neuronal firing rates and MUA activity

Pearson Correlations of firing rates with overall MUA and with MUA time-frequency fluctuations across different frequency bands were calculated, referred to as syncMUA and syncMUADelta, etc.

#### Background noise level

Briefly, it was extracted using an envelope approach, which utilizes the signal envelope distribution to capture the inherent characteristics of the signal noise^41^.

Data from multiple trials were recorded for each subject and the corresponding electrophysiological features were calculated using the median value across all trials of each patient to reduce variability. Furthermore, these features were normalized at the subject level using Z-score normalization. For detailed information on data preprocessing, signal extraction, and feature definition, please refer to our previous work^32^.

### Feature selection

To identify the optimal CT feature combination for individual prognosis, we utilized a machine-learning pipeline composed of five steps (FigureS2A).

#### Step 1 clustering features

Given the redundancy observed among the original 34 CT features (Figure1C, left), we performed hierarchical agglomerative clustering (see ‘Hierarchical agglomerative clustering’ section for details). We set the dendrogram threshold for clustering to ensure all intra-cluster correlation |r| >=0.8^82^. This process yielded 21 clusters with a minimum intra-cluster correlation of r=0.824, ensuring significant homogeneity within each cluster. The relationships among 21 clusters are depicted in FigureS2D, and the features included in each cluster are listed in TableS2. We calculated the mean value within each cluster, designating this average as the cluster-center feature for subsequent analyses to reduce redundancy. Feature values and Importance of Single Feature (Isf) are displayed in FigureS2E.

#### Step 2 filtered feature selection

The filtered feature selection approach assigns relevance scores to each feature based on predefined metrics, selecting only those exceeding a specified threshold. This method is less computationally demanding and suitable for high-dimensional data. Moreover, as it is independent of any machine learning algorithms, filter methods are free from classifier bias, which helps in reducing overfitting^83,84^. For 21 cluster-center features, we assessed their statistical significance, Effect Size, and Fisher Score ranking (details in the ‘Feature importance’ section), respectfully. Criteria for selection included: 1) statistical significance (Wilcoxon Rank Sum Test, p < 0.05), 2) medium to large effect sizes (|r| > 0.3), and 3) the Fisher Scores above a defined inflection point^85^ (FigureS2B).

#### Step 3 embedded feature selection

This approach integrates the selection process directly within the classifier algorithm itself, specifically utilizing a prognosis prediction model equipped with Lasso regularization (details in the ‘Prognosis prediction model’ section). This method inherently penalizes and reduces the weight coefficients of less contributive features, thereby effectively sparsifying feature weights and mitigating overfitting^83,84^. We applied this embedded feature selection technique across four distinct perspectives established in Step 2. These included both the original unfiltered set of 21 cluster-center features and those refined through filtering, based on criteria of statistical significance, effect size, and Fisher Scores.

#### Step 4 feature aggregation

In this step, we combined outputs from the four selection perspectives of Step 3. Features with model weights above 0.05 were aggregated, leading to the selection of clusters No. 2, No. 6, No. 12, No. 15 and No. 18 (TableS2), including six CT features: MUAstabilityTheta (stab-θ), MUApowerTheta (pow-θ), MUApowerAlpha (pow-α), Firing Rates, syncMUAGamma (sync-γ), and MUAstabilityHGamma (stab-hγ) (FigureS2F).

#### Step 5 wrapper feature selection

Wrapper methods evaluate various subsets of features to identify the best-performing combination for a specific classifier algorithm. This approach often yields a smaller set of features with strong discriminative power^83,84^. In our study, we iterated over all possible combinations of the six previously identified features, always including stab-θ (cluster No. 6) due to its significant weight (FigureS2F). This iterative process yielded 32 unique feature combinations, each evaluated for predictive performance using a prognosis prediction model with L2 regularization to optimize accuracy and prevent overfitting (details in the ‘Prognosis prediction model’ section, FigureS2G). These 32 feature combinations were categorized into high accuracy (topAcc) and low accuracy (lowAcc) groups based on their prediction performances, with top accuracy defined as accuracies exceeding 95%. To ensure robustness within topAcc group, we calculated a stable-importance index for each feature, assessing its consistent significance across different combinations within topAcc group (FigureS2G, rightmost column). TableS3 lists each feature’s stable-importance strength along with its average weights in both the topAcc (aveWtop) and lowAcc (aveWlow) groups. Eventually, pow-α and firing rates were excluded from the final optimal feature set (Figure1C, right). This exclusion was likely due to their lower stable-importance strength, minimal aveWlow compared to other features, and their own aveWtop being comparatively lower than their aveWlow.

### Prognosis prediction model

A previous study has measured consciousness states by quantifying arousal and awareness with a deep learning method, specifically using a nested leave-one-participant-out (LOPO) methodology^43^. Similarly, we employed a nested LOPO to evaluate the DoC prognosis prediction performance of the candidate feature subsets. This method utilizes a specialized variant of LOPO cross-validation, implementing a linear SVM as the classifier (FigureS4A).

Specifically, the dataset (N=23) is divided into outer-train and outer-test folds. In each outer fold, all participants except one—the target participant—form the outer-train set (N-1 patients), with the target participant serving as the outer-test set.

For hyper-parameter tuning within each outer-train fold, we utilized LOPO cross-validation combined with a grid search strategy. The training data in the outer-train fold is further subdivided, with N-2 participants forming the inner-train set and the remaining one participant used for inner-validation. The model with optimized parameters that minimize validation error is then tested on the outer-test participant. The mean average accuracy across all folds is calculated as the final accuracy of the prediction model.

We employed distinct regularization strategies during the feature selection phases. In Step 3, Embedded Feature Selection, we utilized Lasso regularization to sparsify model weights:

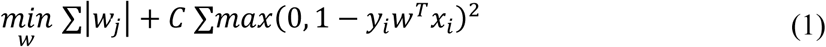

In Step 5, L2 regularization was employed:

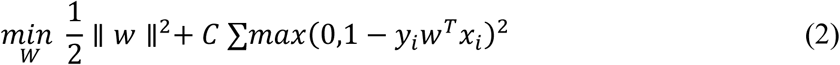

Additionally, to address the sample imbalance—with only 8 CR patients compared to 15 unCR patients—we adjusted the penalty term for misclassifying CR patients to mitigate the impact of this disparity. Our results indicate effective mitigation, as evidenced by the median and mean accuracy distributions for the control group (with shuffled labels) hovering around the chance level (Figure1E).

The LOPO cross-validation procedure enhances the efficient data use and provides unbiased estimates of the classification error across all potential training sets^86^. This method, combined with a linear SVM classifier, not only reduces overfitting but also yields interpretable feature weights, ensuring robust prediction accuracy.

The model’s performance in discriminating between different groups of patients was visualized in the latent features space using both t-SNE and Principal component analysis. T-SNE is a non-linear dimensionality reduction technique that considers relationships between data points and is optimized using Kullback-Leibler divergence. It was implemented using the *’tsne’* function in MATLAB, initialized with the default parameter, *rng(’default’)*.

### Hierarchical agglomerative clustering

Hierarchical agglomerative clustering was performed using an agglomerative approach with the function ‘*clustergram’* in MATLAB. We applied the Weighted Pair Group Method with Arithmetic Mean (WPGMA) as the linkage method, which uses a recursive definition to calculate the distance between two clusters. The distance is computed using weighted average linkage, defined as follows:

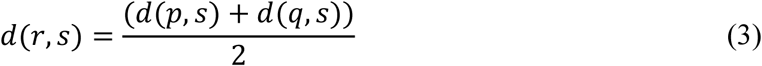

where *r* = *p* ∪ *q* represents the newly formed cluster from clusters *p* and *q*, and *s* is another cluster. We calculated the pairwise distances between CT features using the Spearman Correlation and between patients using the Euclidean distance. Initially, this bottom-up approach treats each data point as an individual cluster. Successive merging of these clusters continues until all data points are unified into a single hierarchical tree (dendrogram), without a predefined number of clusters. The clustering process is finalized by setting a threshold on the linkage strength within the dendrogram to determine the final number of clusters.

### Feature importance

#### Importance of Single-Feature (Isf)

Assessed using the Fisher Score and Effect Size r.

#### Fisher Score

For a feature *i*, the *FisherScore* is computed using the formula^85^:

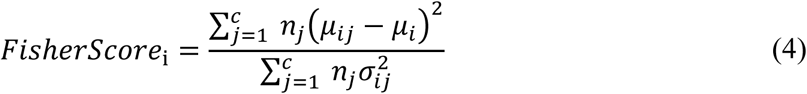

**where** *c* **is the number of classes (CR and unCR),** *n*_*j*_ the number of samples in class *j* **(8 for CR and 15 for unCR),** μ_*i*_ the overall mean of the *i***-th CT feature,** μ_*ij*_ **the mean** within **class** *j*, **and** 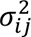 **the variance within class** *j*.

#### Importance in the Multi-Feature Combination (Imf)

Evaluated using the feature weights in the the final predictive model and the Stable-Importance index.

#### Stable-Importance index

This index assesses the consistency of a feature’s significance across different feature combinations within the high accuracy group (FigureS2G). It is defined as the proportion of feature weights exceeding a predefined threshold (≥0.5):

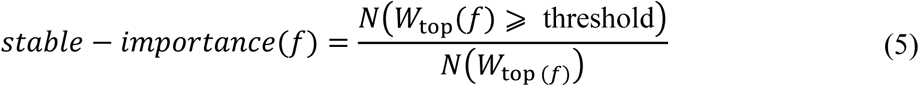

where *W*_top *f*)_ represents the weight of feature *f* in the topAcc group. A higher proportion indicates a feature’s stable significance (TableS3). The choice of threshold does not significantly affect the order of stable-importance strength. For instance, whether the threshold is set between 0.25 and 0.55, or between 0.6 and 1, the top four features with the highest stable-importance indices generally align with the model’s selected features—stab-θ, sync-γ, pow-θ, and stab-hγ (FigureS2H).

### The ablation study

The ablation study was conducted to systematically assess the impact of each feature by removing them one at a time based on their weights. In each round of this process, the prediction model was re-evaluated for prediction accuracy, F1 score, Area Under the Curve (AUC) value of the Receiver Operating Characteristic, Permutation Test p-value, and baseline random accuracy (randAcc), as detailed in TableS4. Initially, the feature stab-hγ was removed due to its minimal weight. The model was then retrained with the remaining features. This iterative process was repeated until only the two most significant features, stab-θ and pow-θ, remained, with stab-θ ultimately emerging as the top-ranked feature.

#### F1 score

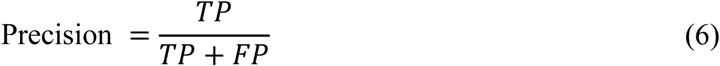

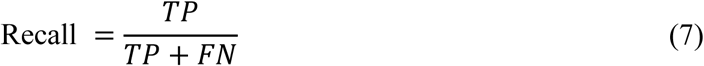

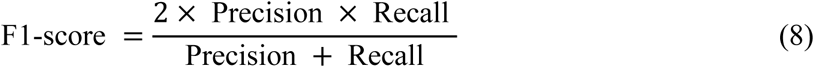

where true/false positive predictions (*TP/FP*) and true/false negative predictions (*TN/FN*) are used. This metric combines precision and recall to provide a single score that balances both the false positives and false negatives.

#### Baseline Random Accuracy (randACC)

Estimated from the mean accuracy of 1000 shuffled iterations in a permutation test, this measure serves as a control. It represents the expected accuracy from a random binary classification, where each permuted dataset undergoes the same prediction analysis as the original dataset.

### CT metric for DoC prognosis

We developed a metric based solely on CT characteristics, using a weighted average of the selected feature weight coefficients (FigureS4A). The formula for the *CT metric* is:

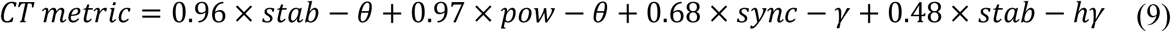

To evaluate the efficacy of this metric, we compared the prediction scores of each patient obtained from the final optimal model with those derived directly from this CT metric (FigureS4B). Our findings demonstrate that the CT metric reliably represents the outcomes predicted by the final optimal model, as evidenced by: 1) the consistency of correct predictions for each patient by both the CT metric and the model, and 2) the minimal discrepancies between their respective prediction scores.

### Subject similarity

Subject similarity was quantified using the normalized reciprocal of the Euclidean distance between each pair of patients. Specifically, for two patients, *P*1 and *P*2, each characterized by (*F*1, *F*2, …, *FN*), the similarity *s* is calculated as follows:

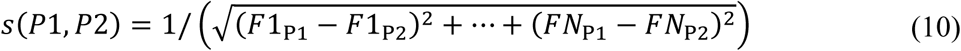

This similarity score *s* was then subjected to min-max normalization across all patient pairs within the subject-similarity matrix to standardize the values. We applied this similarity metric under various conditions, comparing within and between different patient groups (CR/unCR, Figures2F and S5B), and among subgroups (Ⅰ/Ⅱ/Ⅲ, FigureS6B).

### CT-Based subgroups of DoC identification

#### Identifying DoC subgroups through hierarchical agglomerative clustering

This methodology, previously leveraged in anesthesia research, has revealed novel patterns of desynchronized neuronal activity, challenging traditionally presumed monotonic correlations between anesthesia depth and neuronal activity^37,87^. Motivated by these findings, we employed the same hierarchical agglomerative clustering approach to delineate CT-based subgroups of DoC, utilizing Euclidean distances between patients’ CT features. To determine the optimal number of clusters, we applied the ‘elbow’ method^37^. Specifically, we plotted the within-cluster sum of distances, which were min-max normalized, against the number of clusters (FigureS6A). The point of greatest curvature, the ‘elbow,’ indicative of the sharpest shift in distance reduction, was identified by the maximum of the second-order difference of the distance-K curve.

In our result, we identified distinct CT activity patterns within DoC patients. Based on the ‘elbow’ method, we selected K=4, resulting in the formation of four distinct clusters. Subgroups Ⅰ and Ⅱ exhibited clustering (Figures3A and S6B). The remaining two clusters included one cluster with only Patient No. 6. Due to their similarities in CT profiles, and characterized by elevated stab-θ strength (Figures3A and S6C), these clusters were merged into a single entity, Subgroup Ⅲ.

#### Venn diagram analysis of feature combinations

For each patient, a feature was categorized as high-value if its value exceeded the median value of the cohort. We analyzed the distribution of these high-value feature within each DoC group using Venn diagrams (Figures2G, 2H, and 3C-3E). To address potential sample imbalances among the DoC groups (CR/unCR or Subgroups Ⅰ/Ⅱ/Ⅲ), we implemented oversampling with replacement to balance the classes. This was achieved by resampling each class to match the size of the largest class, and then employing bootstrap resampling 1,000 times to accurately obtain the distribution of CT features among these groups^88^. The averaged results from these resampling iterations were used to construct the final Venn diagrams. Additionally, the accompanying pie chart illustrates the distribution of patients based on the number of simultaneously high-value features.

### Biophysical simulation model

#### Model overview

Our model simulated interactions between inhibitory neurons within the TRN and excitatory neurons in the thalamocortical relay nuclei, both of which were crucial in modulating consciousness and arousal^21,22,45,50–53^. The excitatory neurons were further categorized into matrix and core populations within the CT and SR, respectively. Each nucleus in our simulation consisted of 100 neurons.

#### Input and connectivity

Each neuron received random excitatory input modeled by a Poisson spike train with a base discharge probability P of 0.5 for CT/SR and 0.15 for TRN^54^. Connectivity was established with one-to-one connections between SP and TRN and one-to-many connections from CT to TRN at a probability of 0.15^89^. Additional connectivity details are provided in TableS5.

#### Neuron dynamics

Neuron dynamics were modeled using the integrate-and-fire-or-burst approach^90^, adapting the classical conductance-based leaky integrate-and-fire model to include low-threshold bursting (LTB) (Eq. 15-17). This adaptation was critical for simulating thalamic discharge patterns under deafferentation conditions, essential for understanding thalamocortical rhythmic activities. Parameters related to thalamic neuronal dynamics are outlined in TableS6.

#### Model equations

The membrane potential *V* of neurons is modelled by:

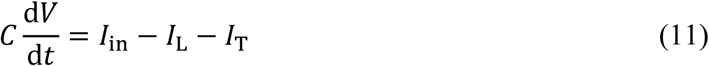

where *C* is the membrane capacitance, *I*_in_ represents the synaptic input current, *I*_L_ is the leak current, and *I*_T_ is the total calcium current. Neurons spike when *V* exceeds a threshold *<u>V</u>*_θ_, after which it is reset to *V*_r*eset*_.

The synaptic input current *I*_in_ is derived from the sum of excitatory and inhibitory synaptic currents:

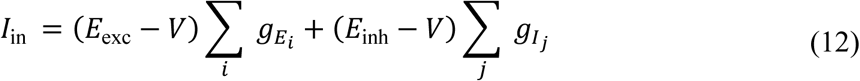

where *E*_exc_ and *E*_inh_ represent the excitatory and inhibitory reversal potentials, respectively, and *g*_*Ei*_ and *g*_*Ii*_ are the conductances for excitatory and inhibitory synapses.

Synaptic conductance decays exponentially with a time constant τ_s_, modulated by presynaptic firing events after a connection-specific delay. Specifically, when a presynaptic neuron fires, it causes a temporary increase in synaptic conductance by *g*_*E*/*I*_, which then follows an exponential decay:

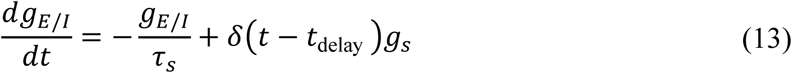

In this equation, δ(*t* − *t*_delay_) represents a delta function that activates at the delay time *t*_delay_ following presynaptic firing. The leak current *I*_L_is defined by:

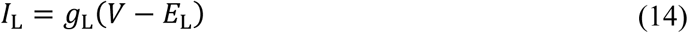

with *g*_L_ as the leak conductance and *E*_L_ the leak reversal potential.

As DoC is associated with a reduced neuronal resting membrane potential (RMP) due to potassium leakage currents^8,55,56^, a lower RMP is set for simulating DoC conditions. This RMP is lower than both the healthy RMP and the calcium LTB threshold.

The calcium current, *I*_T_, is critically dependent on the inactivation state h and is governed by the equation:

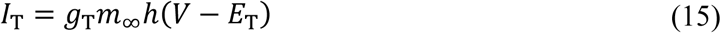

Here, *g*_T_ is the calcium conductance, *E*_T_ is the calcium reversal potential, and m_∞_ is defined by the heaviside step function H, as follows:

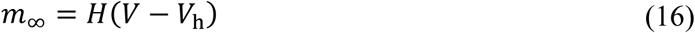

This function ensures that activation occurs instantaneously above a threshold voltage _h_. The inactivation variable ℎ dynamically adjusts based on the membrane potential *V*, engaging in a two-stage de-inactivation–activation process. This process allows ℎ to approach zero when *V* exceeds _h_ (depolarized conditions) and to approach one under hyperpolarized conditions, with transitions governed by specific time constants 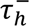 and 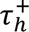:

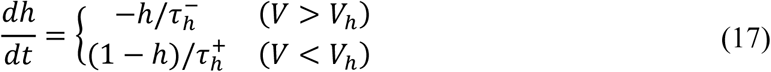

#### Modeling various afferent conditions

Varied levels of structural and functional deafferentation for both CT and SR were modeled through different afferent conditions, encompassing three main factors:

***1) Proportion of deafferented neurons (deaffRatio)***: This parameter represents the percentage of neurons undergoing deafferentation, with values ranging from 0% to 100%. If there is no residual input left, i.e., complete deafferentation, it leads to an absence of discharging activity, simulating conditions akin to neuronal damage *per se* within the network.
***2) Afferent magnitude (affMag) of residual input***: This factor adjusts the magnitude of residual input preserved in neurons that have undergone partial reafferentiation (Figure5A). The afferent magnitude (affMag) varies from 0% to 100% relative to normal spike train input conditions (discharge probability P of 0.5 for the CT neurons).
***3) Afferent stability (affStab) of residual input***: Stability is manipulated by introducing “silent periods” that intermittently suppress the input spike train (Figure5A). This approach models alternating phases of normal activity and “silent periods” for all neurons to simulate intermittent spiking inputs under slow oscillation and/or burst suppression, which can significantly alter the overall membrane potentials across CT neurons. The impact of these “silent periods” on input stability is quantified from 0% (complete suppression during “silent periods”) to 100% (full preservation of spikes during “silent periods”). For instance, an affStab of 50% means that during “silent periods”, half of the expected input spikes are lost compared to normal conditions.

#### Four stages of consciousness

Our simulations delineated four distinct stages of consciousness based on various afferent conditions:

##### Stage I

Most CT neurons (deaffRatio ≥ 90%) are under complete deafferentation, exhibiting a quiescent state.

##### Stage II

Partial restoration in CT neurons (deaffRatio = 30%) leads to slightly higher firing rates than Stage Ⅰ but remains insufficient to produce theta rhythms.

##### Stage III

CT neurons undergoing reafferentiation reach a theta rhythmic state, characterized by affMag of 4%, which induces the largest amplitude of theta activity observed.

##### Stage IV

All CT neurons are fully restored to a healthy profile, with complete reafferentiation (deaffRatio = 0% and affMag = 100%), and normal membrane potential restored to the normal RMP.

#### Analysis of simulated CT activities

To analyze the simulated CT activities, we first calculated the firing rates by averaging the number of spikes per second across all CT neurons. Subsequently, we averaged the membrane potentials of all CT neurons at each timestep, beginning three seconds post-initiation. This averaged data provided the foundation for further analysis, including:

##### Power and stability assessment

We evaluated pow-θ and stab-θ using methodologies consistent with those applied to empirical MUA signals, including normalized power and signal stability for MUA activity in the theta band (details in the ‘Signal processing and feature extraction’ section). Notably, compared to LFP—a proxy for synaptic input over an extended range—MUA provides more localized information regarding population outputs^33,34^. Thus, in this context, the simulation model that describes the rhythmic activity of CT outputs (small size nuclei) more closely aligns with the rhythms observed in MUA.

##### Power spectral density (PSD)

The PSD of CT activities was calculated to capture frequency components ranging from 0 to 80 Hz.

### Statistical analysis

#### Wilcoxon Rank Sum Test

We employed the Wilcoxon Rank Sum Test (two-tailed, unpaired) to identify significant differences between the CR and unCR groups. This non-parametric test evaluates the median differences between two independent samples.

**Effect Size r** for Wilcoxon Rank Sum Test. The r is calculated as

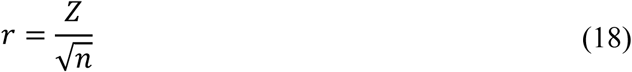

where n is the total number of observations. The effect size is categorized as small (0.1∼0.3), medium (0.3∼0.5), and large (>0.5)^91^.

#### Spearman Correlation

It was applied to assess the correlation relationships between CT features in hierarchical agglomerative clustering. It evaluates the monotonic relationship between variables, utilizing the ranks of the data.

#### Pearson Correlation

Used to calculate the synchronization between neuronal activity and MUA, this method assesses the linear relationship between two continuous variables.

#### Permutation Test

The Permutation Test was utilized to validate the robustness of our model’s accuracy against random chance. To generate a null distribution, we randomly shuffled the labels (unCR and CR) 1,000 times to create 1,000 simulated datasets. Each dataset was then analyzed using the same prediction model that was applied to the original data. Model performance ranking within the top 5% of this null distribution was considered statistically significant, with a p-value of less than 0.05.

#### Fisher’s Exact Test

To evaluate the statistical independence between two categorical variables in small datasets, we employed Fisher’s Exact Test. This test is particularly useful when expected frequencies in any of the cells of a contingency table are below 5. For scenarios involving three categories, we utilized an extended version of Fisher’s Exact Test (comparisons among three etiologies and three subgroups). This adaptation accommodates larger tables and a broader range of categorical variables. The extended test employs a network algorithm for exact computation of p-values, effectively handling the complexities of larger contingency tables.

#### Odds Ratio (OR)

Implemented by the conditional maximum likelihood estimate, the Odds Ratio quantifies the strength and direction of the association between two categorical variables. This approach is particularly suited for small sample sizes or unevenly distributed categorical data, ensuring robust and unbiased estimates of association strength. An OR greater than 1 indicates a positive association, less than 1 indicates a negative association, and equal to 1 indicates no association.

#### Effect Size (Cohen’s h)

To quantify the magnitude of association between the two categorical variables in our analysis, we calculated Cohen’s h, an effect size used to measure the difference between two proportions. It is defined as:

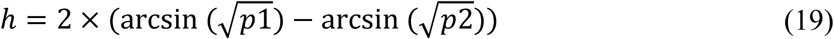

where *p*1 and *p*2 are the proportions from two independent groups.

#### Kruskal–Wallis (KW) Test

The KW test to statistically assess differences in CT features among various etiologies, followed by a post-hoc analysis using Conover’s All-Pairs Rank Comparison Test.

#### Effect Size (η^2^)

The effect size for KW was η^292^ who was defined as

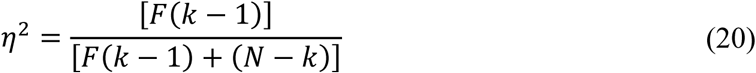

where *N* was the sample size, *k* was the group number and *F*(*k* − 1) was the statistics *F* with *k* − 1 degree of freedom.

#### Significance level

Significance levels are represented as follows: no significance (n.s.) for p ≥ 0.05; * for p < 0.05; ** for p < 0.01; *** for p < 0.001.

All data preprocessing and analyses were conducted using MATLAB R2023 (MathWorks, Inc.) and RStudio (R 4.3.2). Fast Fourier Transforms were performed using the *’pwelch’* function in MATLAB. Spearman Correlation and Pearson Correlation were computed using the *’corr’* function in MATLAB.

The Wilcoxon Rank Sum Test was implemented using the *’wilcox.test’* package, and its effect size (r) was computed using the *’qnorm’* package to calculate the Z statistic in R. Fisher’s Exact Test (both the standard and extended tests) and Odds Ratio calculations were performed using the *’fisher.test’* package in R. The KW test was implemented using the *’ kruskalTest’* package, and its post-hoc analysis (Conover’s All-Pairs Rank Comparison Test) was conducted with the *’kwAllPairsConoverTest’* package in R.

## Acknowledgments

This study was supported by the National Natural Science Foundation of China (82272118), the National Key Research and Development Program (2023YFB4706100), and the Beijing Natural Science Foundation (7232046). S.L., Research Director at the Belgian National Fund for Scientific Research (FRS-FNRS), is supported by the European Foundation of Biomedical Research FERB Onlus, the Generet fund of the King Baudouin Foundation, and the Mind Care International Foundation. We acknowledge Binghao Yang and Shenghao Cao for technical help, and Ruoshui Xu for helpful discussions.

## Author contributions

Conceptualization: S.Y., J.H.H., H.R.Z., and S.L.

Resources: J.H.H., Q.Q.G., Y.Y.D., Y.T.Z., and L.X.

Methodology: S.Y., H.R.Z., S.W., and X.L.

Investigation: S.Y., J.H.H., H.R.Z., S.L., Q.Q.G., S.W., X.L., Y.Y.D., Y.T.Z., and L.X.

Visualization: S.Y., H.R.Z., and S.L.

Funding acquisition: S.Y., J.H.H., and S.L.

Project administration: J.H.H. and S.Y.

Supervision: S.Y. and J.H.H.

Writing – original draft: S.Y., H.R.Z., J.H.H., and Q.Q.G.

Writing – review & editing: S.Y., H.R.Z., S.Y., J.H.H., and Q.Q.G.

## Data availability

All data and methods are available in the main text file.

Datasets supporting the findings of this study can be obtained from the corresponding authors upon reasonable request.

---

**FigureS1.**
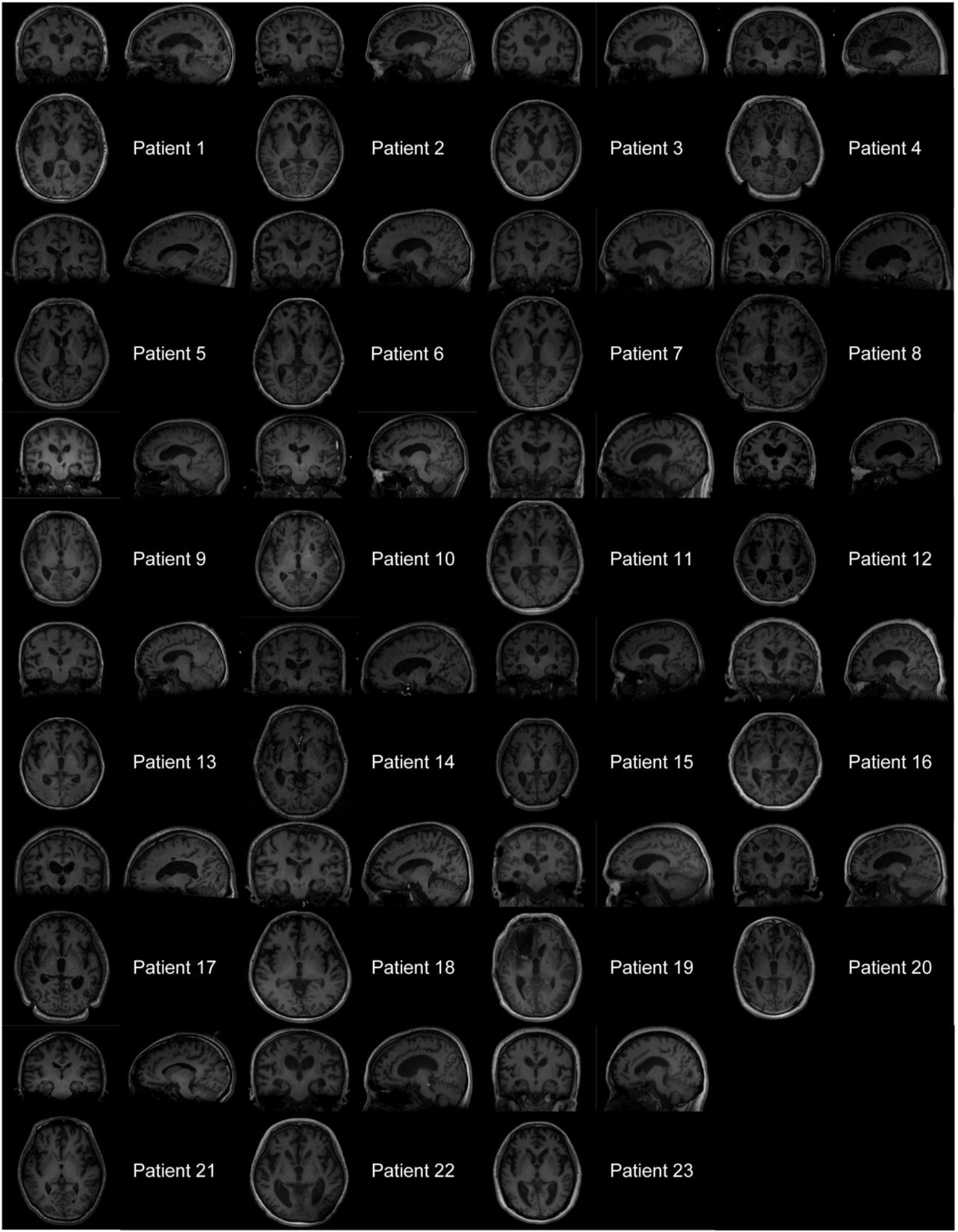
Preoperative MRI scans of all DoC patients. Typical sections-coronal (top left), sagittal (top right), and axial (bottom left)-of thalamus are shown for each patient in each panel. These MRIs illustrate relatively well-preserved brain structures across all patients.

**FigureS2.**
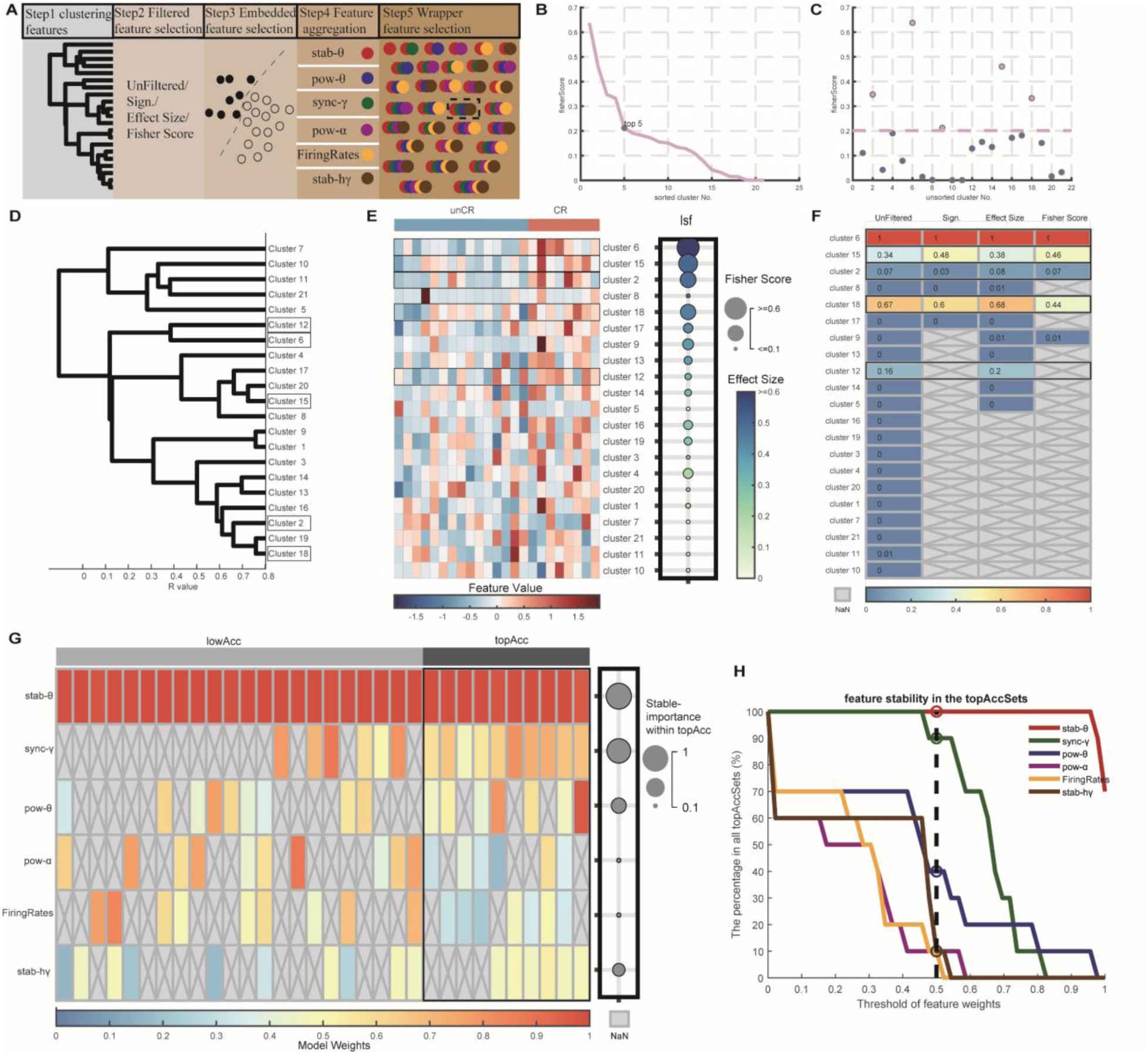
Feature selection pipeline for DoC prognosis. **(A)** Overview of the five sequential stages of the feature selection pipeline: ‘Unfiltered’, ‘Sign.’, ‘Effect Size’, and ‘Fisher Score’ refer to statistical significance, effect size, and Fisher Scores. In Step 5, the final selected four features of the CT metric are highlighted with dashed borders. **(B)** Identifies the inflection point for 21 cluster-center features sorted by Fisher Scores, highlighting the top 5 as having higher values. **(C)** The original Fisher Score distribution for the datasets corresponding to those in (**B**). The red dotted line represents the filtering threshold for Fisher Scores. **(D)** Hierarchical agglomerative clustering of the 21 clusters (Step 1). Clusters No. 2 (Firing Rates), No. 6 (stab-θ), No. 12 (stab-hγ), No. 15 (pow-θ and pow-α), and No. 18 (sync-γ) are highlighted with black borders. **(E)** Electrophysiological profiles of 21 cluster-center features. Heatmap across patients showing higher values in CR (n=8) compared to unCR (n=15). The importance of Single-Feature (Isf) is indicated by the Fisher Score (circle size) and effect size (color intensity). **(F)** Summary of feature weights obtained through embedded feature selection (Step3) from the four selection perspectives, including ‘Unfiltered’, and filtering through ‘Sign.’, ‘Effect Size’, and ‘Fisher Score’ (defined in Step 2). Features with model weights above 0.05 (Cluster No. 2, 6, 12, 15, and 18) were highlighted with black borders and aggregated for subsequent analysis (Step 4). **(G)** 32 iterated feature combinations in the wrapper feature selection (Step 5) were divided into high accuracy (topAcc, accuracy exceeding 95%, marked with black borders) and low accuracy (lowAcc) groups. The consistent significance across combinations within topAcc (circle size) is shown with Stable-Importance strength for each feature. **(H)** Threshold Curve for Stable-Importance index: Depicts the stability of selected features as a function of weight thresholds. It measures the possibility of each feature weighing above different thresholds in the topAcc combinations.

**FigureS3.**
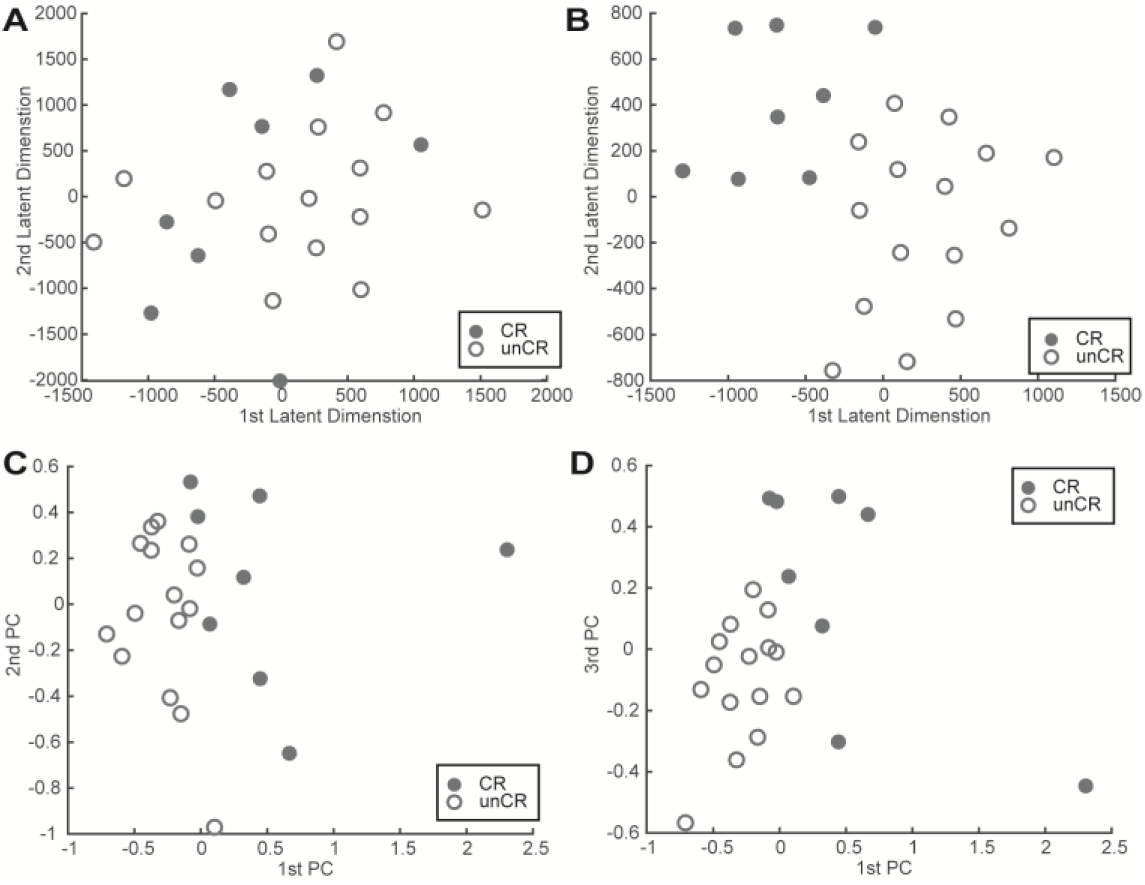
Distribution of patients in feature space. **(A and B)** T-SNE analysis before and after feature selection. The scatter plot displays substantial overlap before feature selection (**A**) and shows improved discrimination after selecting four key features (**B**) in the t-SNE latent feature space. **(C and D)** Principal component analysis post-feature selection: The improved discrimination between patient groups is also evident in the 1^st^ principal component (PC) compared to the 2^nd^ / 3^rd^ PC after feature selection.

**FigureS4.**
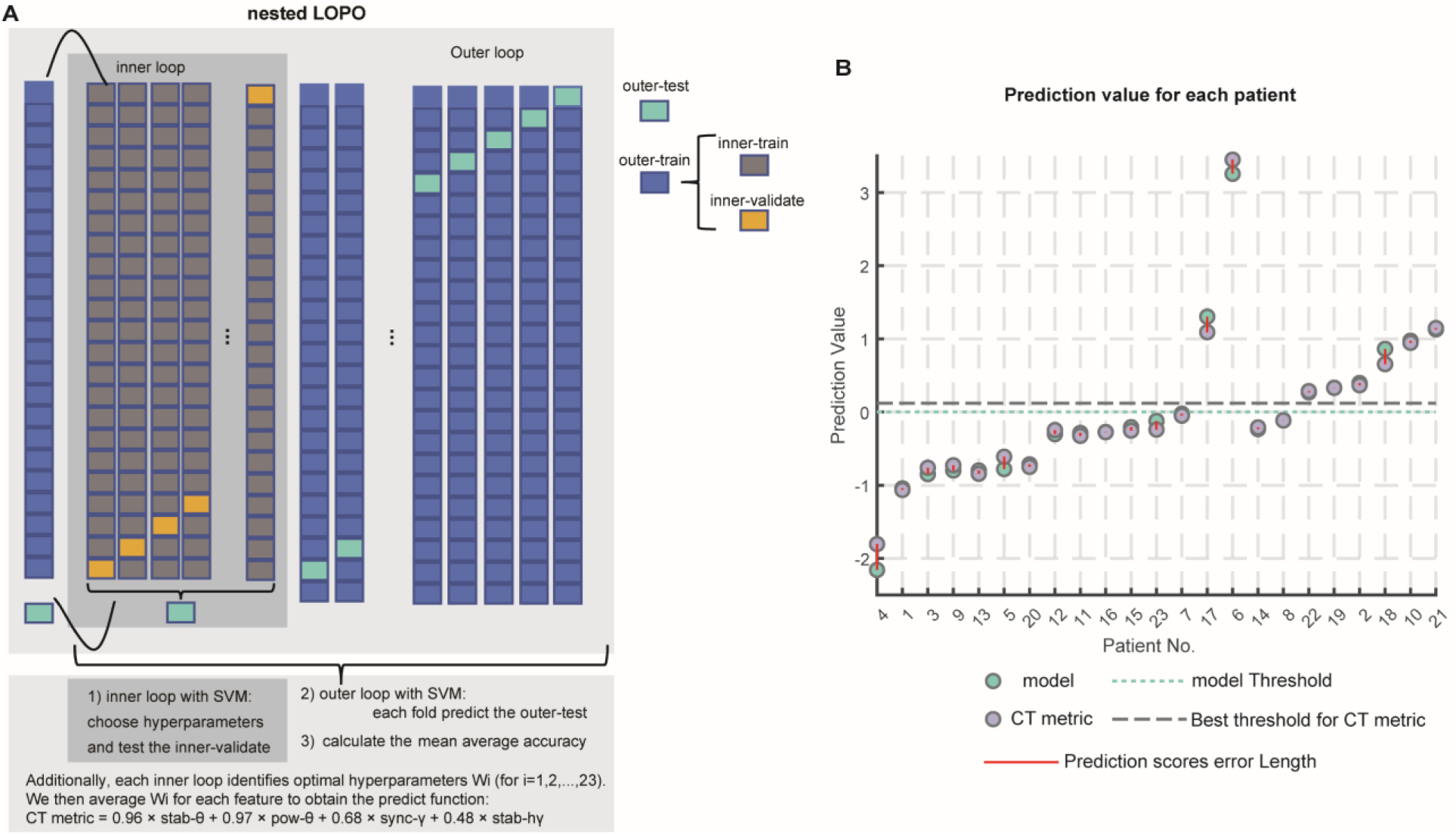
Nested LOPO cross-validation and prognosis prediction using the CT metric. **(A)** Diagram illustrating the nested leave-one-participant-out (LOPO) cross-validation process using a linear SVM classifier. The process is depicted with both inner and outer loops; the inner loop is designated for hyperparameter tuning and validation, while the outer loop is utilized for final testing. **(B)** The prognosis prediction values of the prediction model and CT metric for each patient. The CT metric is plotted alongside actual model predictions, illustrating the alignment between the two.

**FigureS5.**
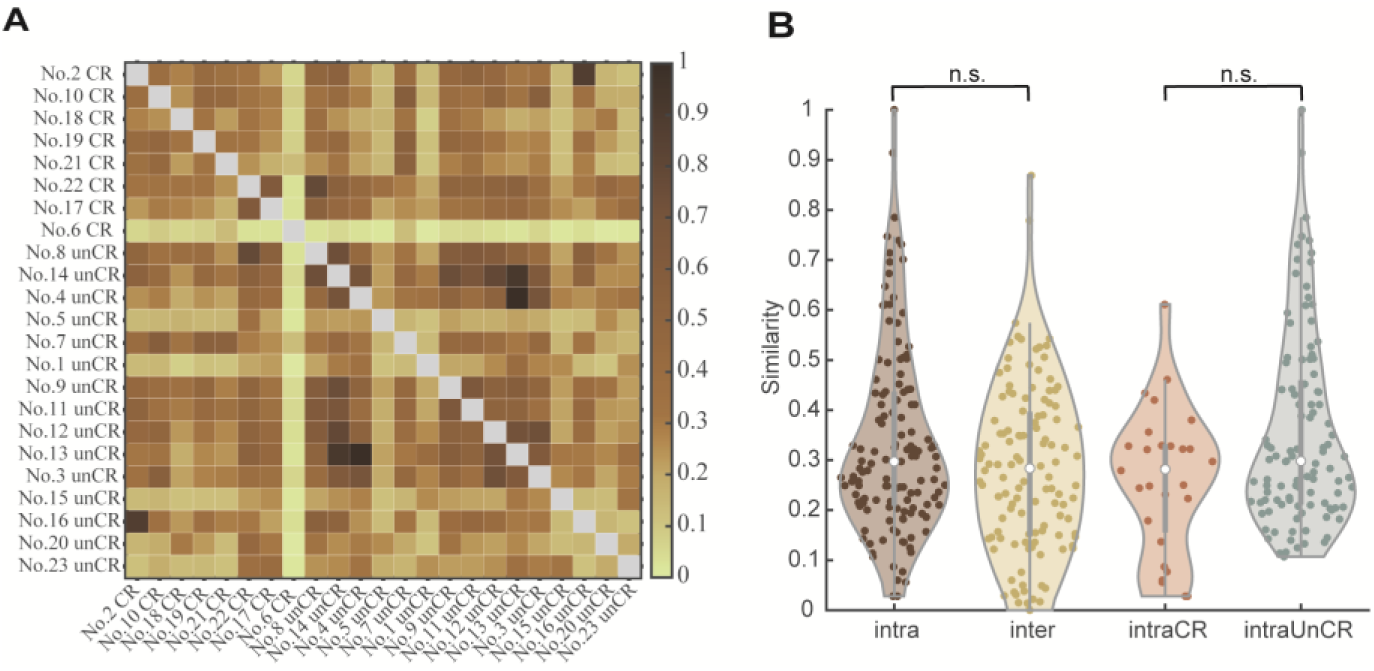
Similarity analysis of CT features among patients before feature selection. **(A)** Heatmap of pairwise similarities between patients, categorized into CR and unCR groups. **(B)** Violin plots show the distribution of similarity within (’Intra’) and between (’Inter’) groups.

**FigureS6.**
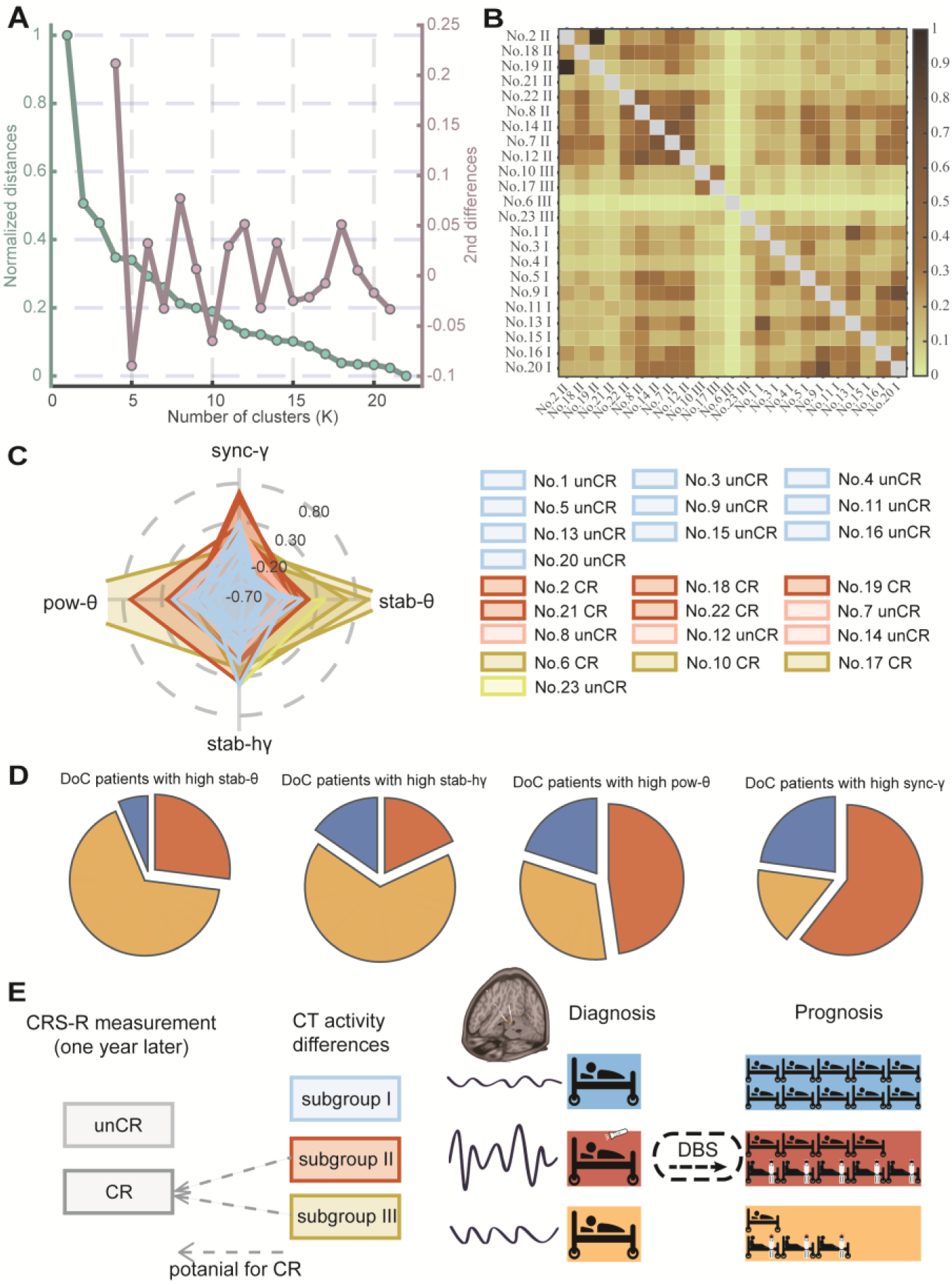
Subgroup identification in DoC and Their CT-Activity Profiles. **(A)** Dendrogram from hierarchical agglomerative clustering. The elbow method suggests that the optimal number of clusters is four (k=4), indicating distinct subgroup formations based on CT features. **(B)** Heatmap of pairwise similarities between patients, grouped into three subgroups. **(C)** Radar Chart illustrating distributions of CT features across patients, color-coded by the three identified subgroups. **(D)** Distribution of four CT-Activity features across subgroups: Shows that higher-value pow-θ and sync-γ are primarily found in Subgroup Ⅱ, while higher-value stab-θ and stab-hγ are mainly present in Subgroup Ⅲ. **(E)** Behavioral measurements (diagnosis and prognosis) of three subgroups: Patients in the CR group are differentiated into Subgroups Ⅱ and Ⅲ based on their distinct CT profiles. Subgroup Ⅰ, characterized by subdued CT activity, consists only of UWS patients who remained unCR post-treatment; Subgroup Ⅱ, with pronounced theta rhythms, consists mainly of MCS-patients, over half of whom improved to CR status post-treatment; Subgroup Ⅲ, with stable theta rhythms, predominantly includes UWS patients, most of whom are categorized as CR in prognosis.

**FigureS7.**
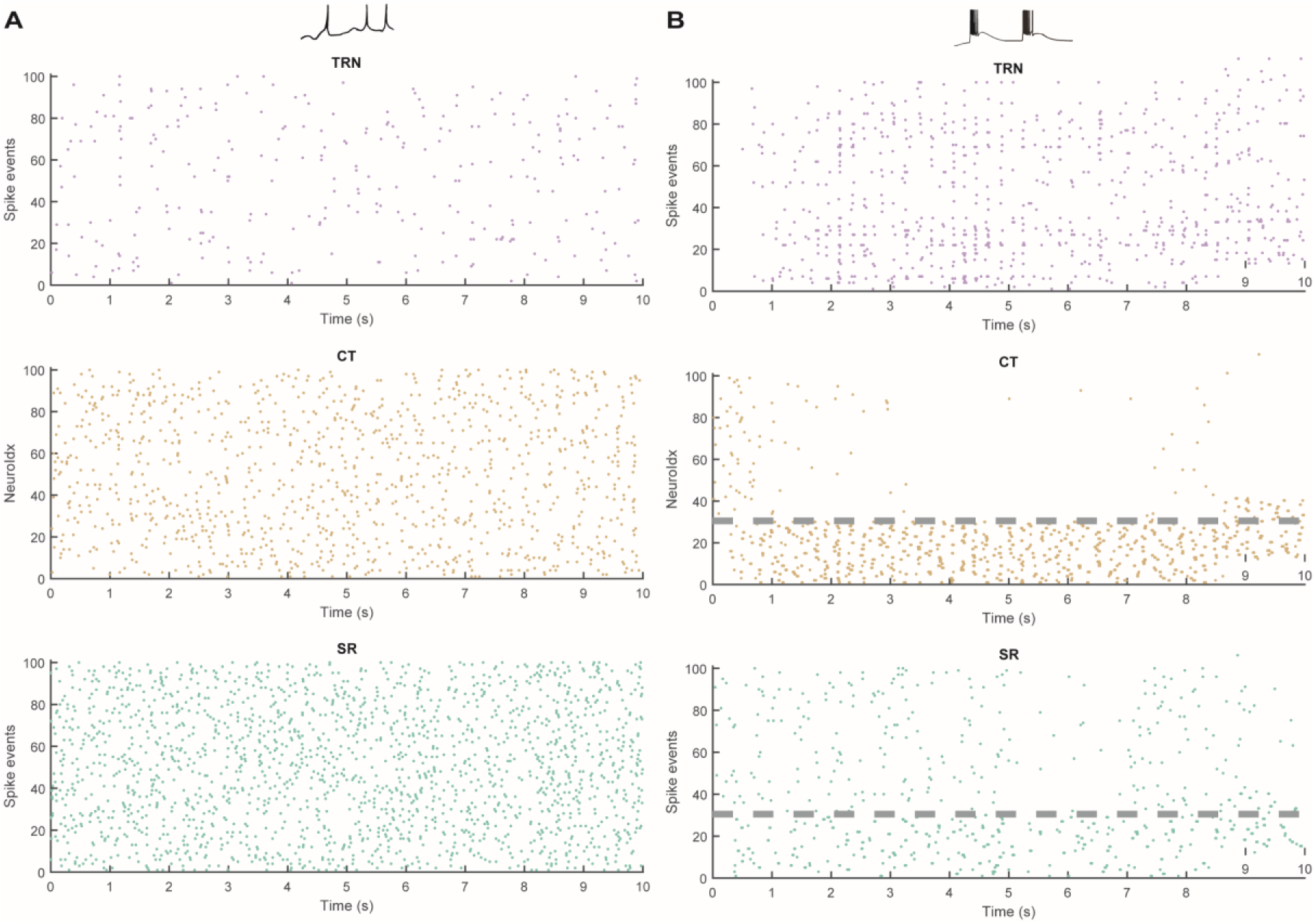
Raster plots for tonic and bursting regimes in thalamic neurons. **(A)** Tonic regime: Activity under normal conditions with a membrane potential (E_L_ = -65 mV) and typical afferent inputs. Raster plots show the spiking behavior over 10 seconds for neurons in the thalamic reticular nucleus (TRN), corticothalamic (CT), and specific relay nuclei (SR). **(B)** Bursting regime: Activity under a lower resting membrane potential (E_L_ = -67 mV) and deafferent inputs, with 30% deafferented neurons (deaffRatio) and 4% of the magnitude of residual afferent input (affMag).

**FigureS8.**
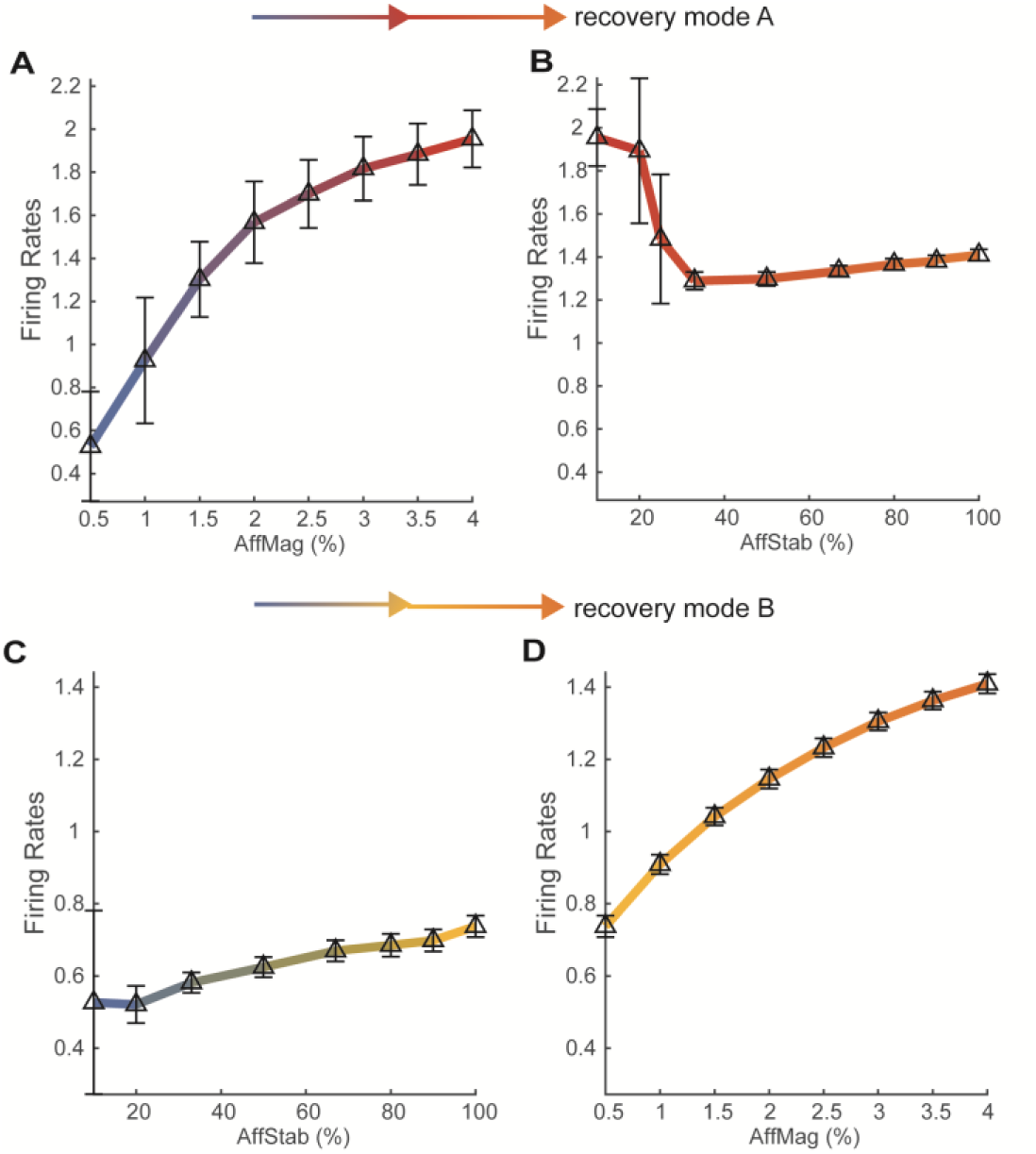
Modulation of firing rates by afferent stability and magnitude. **(A andB)** Recovery Mode A: (**A**) Demonstrates a substantial rise in firing rates with increasing afferent magnitude (AffMag). (**B**) Reveals minimal impact on firing rates, as afferent stability (AffStab) increases. **(C and D)** Recovery Mode B: (**C**) Exhibits slight changes in firing rates with varying afferent stability (AffStab). (**D**) Shows a significant monotonic increase in firing rates with increasing afferent magnitude (AffMag).

**FigureS9.**
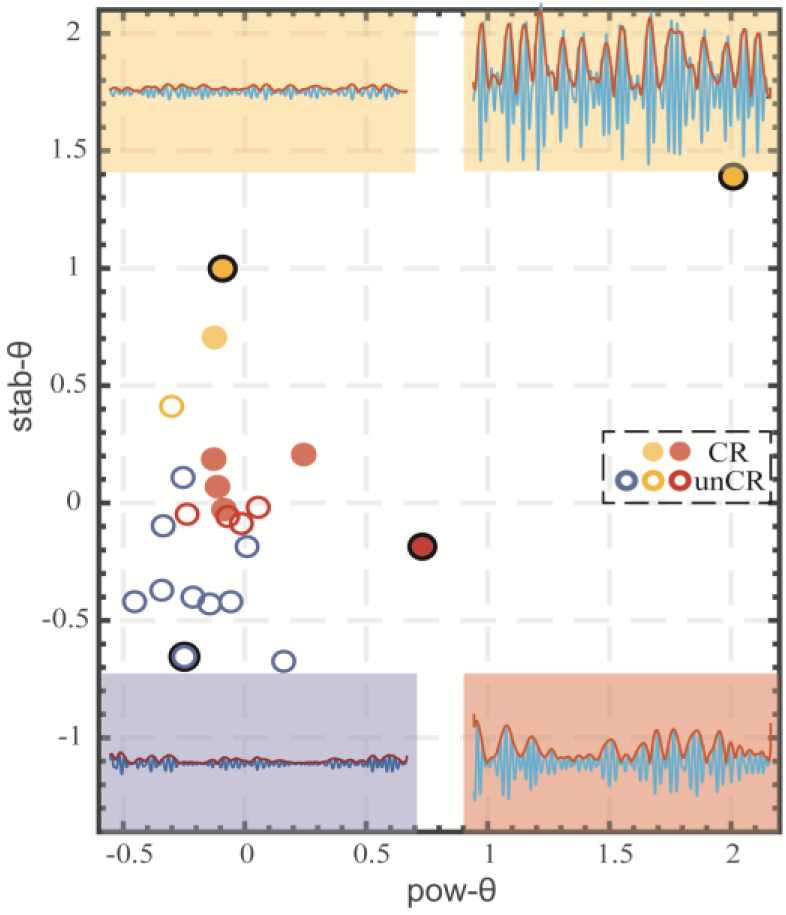
Empirical distribution of patients from three subgroups. This figure showcases the empirical data across three subgroups, each characterized by distinct patterns in the power (pow-θ) and stability (stab-θ) of theta rhythms: Subgroup Ⅰ (blue) consistently displays lower values in both pow-θ and stab-θ. Subgroup Ⅱ (red) is characterized by high pow-θ but low stab-θ. Subgroup Ⅲ (yellow) typically shows high stab-θ but low pow-θ, with an outlier in the upper right corner exhibiting high values in both features. The insets provide detailed theta band profiles in the CT from four example patients: Patient No. 4 in blue with low pow-θ and stab-θ; Patient No. 21 in red with high pow-θ and low stab-θ; Patient No. 17 in yellow (top left) with high stab-θ and low pow-θ; and Patient No. 6 in yellow (top right) showing high levels in both features.

**FigureS10.**
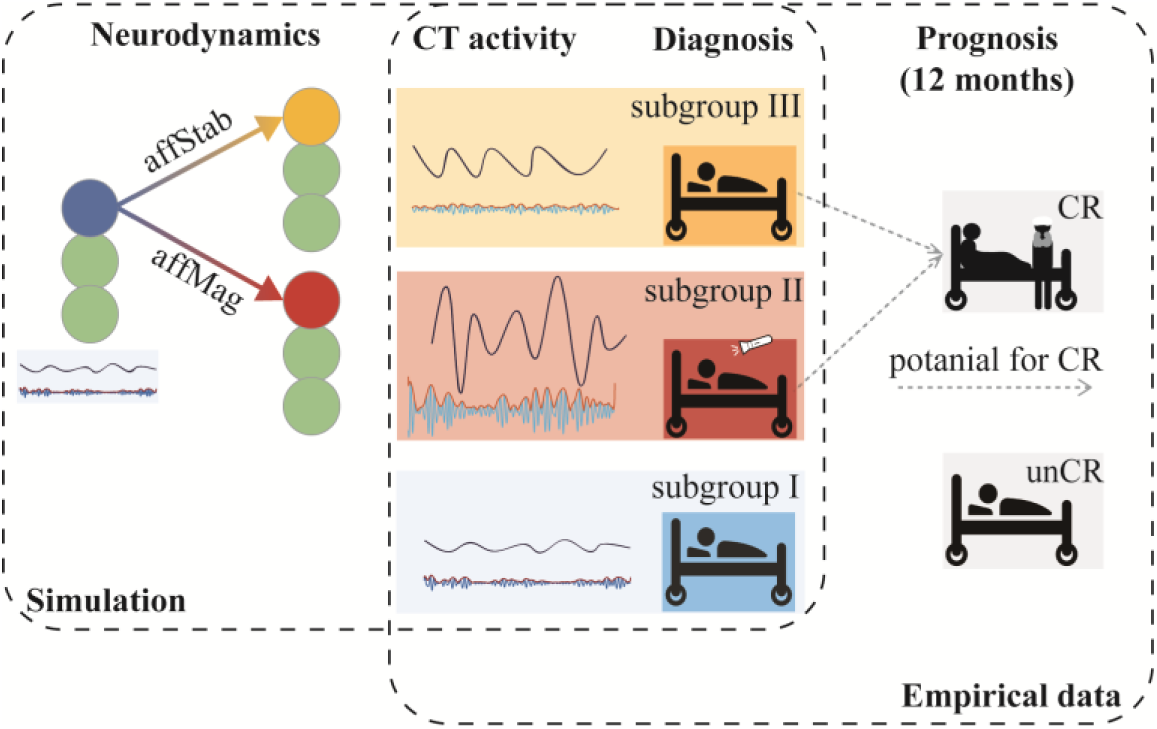
Schematic Based on empirical data of CT activity, we identified two subgroups (II and III) with potential for consciousness recovery. These subgroups exhibit distinct electrophysiological patterns in the CT: Subgroup III is characterized by stable theta band activity, whereas Subgroup II displays greater amplitude in theta activity. Moreover, these subgroups have different clinical baselines: most MCS patients belong to Subgroup II, for example, demonstrating visual pursuit in response to a flash. In contrast, all patients who recovered from UWS were from Subgroup III. Subgroup I shows quiescent CT activity and lacks recovery potential. Additionally, using a simulation model, we explored the neurodynamics underlying these distinct theta rhythm behaviors. This revealed that the stability and magnitude of afferent neuronal input respectively facilitate the transition from silent CT to either stable or high-amplitude theta rhythms.

**TableS1.** Patients’ clinical information.

| Subgroup | Patient number | Gender | Etiology | Age (yrs) | Duration (months) | Preoperative Diagnosis | Preoperative CRS-R | Follow-up Diagnosis | Follow-up CRS-R |
| --- | --- | --- | --- | --- | --- | --- | --- | --- | --- |
| I | 1 | F | Anoxia | 64 | 3 | UWS | 6 (111102) | UWS | 6 (111102) |
| I | 3 | M | Anoxia | 47 | 1.5 | UWS | 6 (102102) | MCS- | 9 (213102) |
| I | 4 | F | Anoxia | 60 | 4 | UWS | 6 (102102) | UWS | 8 (122102) |
| I | 5 | M | BSH | 69 | 4 | UWS | 2 (002000) | UWS | 4 (102100) |
| I | 9 | F | Trauma | 53 | 3 | UWS | 7 (112102) | UWS | 7 (112102) |
| I | 11 | M | BSH | 49 | 5 | UWS | 6 (102102) | UWS | 5 (002102) |
| I | 13 | F | Anoxia | 43 | 3 | UWS | 7 (112102) | MCS- | 12 (233202) |
| I | 15 | M | Anoxia | 41 | 4.5 | UWS | 8 (112202) | UWS | 8 (112202) |
| I | 16 | F | BSH | 64 | 3 | UWS | 3 (002100) | UWS | 4 (102100) |
| I | 20 | M | Trauma | 30 | 3.5 | UWS | 5 (102101) | MCS- | 9 (213102) |
| II | 2 | M | Trauma | 48 | 11 | MCS- | 14 (245102) | eMCS | 20 (445322) |
| II | 7 | M | BSH | 39 | 6 | UWS | 4 (001102) | UWS | 4 (001102) |
| II | 8 | M | BSH | 52 | 6.5 | MCS- | 9 (132102) | MCS- | 9 (132102) |
| II | 12 | M | Anoxia | 59 | 4 | UWS | 5 (002102) | UWS | 6 (102102) |
| II | 14 | F | BSH | 59 | 3 | MCS- | 9 (132102) | MCS- | 8 (032102) |
| II | 18 | M | BSH | 35 | 4.5 | MCS- | 9 (132102) | MCS+ | 19 (455113) |
| II | 19 | M | Trauma | 26 | 9 | MCS- | 9 (132102) | eMCS | 23 (456323) |
| II | 21 | F | Anoxia | 25 | 13 | MCS- | 8 (113102) | MCS+ | 21 (455313) |
| II | 22 | F | Anoxia | 11 | 7 | MCS- | 8 (113102) | eMCS | 14 (333122) |
| III | 6 | M | BSH | 59 | 3.5 | UWS | 4 (012001) | MCS+ | 11 (332102) |
| III | 10 | M | Trauma | 26 | 3 | MCS- | 10 (232102) | eMCS | 23 (456323) |
| III | 17 | M | Trauma | 39 | 3.5 | UWS | 7 (112102) | MCS+ | 12 (333102) |
| III | 23 | F | Anoxia | 43 | 1 | UWS | 7 (112102) | UWS | 7 (112102) |

F: Female; M: Male; BSH: brainstem hemorrhage; Coma: chronic coma; UWS: Unresponsive
Wakefulness Syndrome; MCS-/MCS+: Minimally Conscious State minus/plus; eMCS:
emergence from Minimally Conscious State; CRS-R: JFK Coma Recovery Scale-Revised.

**TableS2.** CT-activity feature clusters and associated features.

| Cluster No. | CT Features |  |  |
| --- | --- | --- | --- |
| 1 | NoiseLevel | MUAMean |  |
| **2** | **FiringRates** |  |  |
| 3 | Spike-clusteringIndex | Spike-clusteringRatio |  |
| 4 | MUAstabilityDelta |  |  |
| 5 | PauseIndex | PauseRatio |  |
| **6** | **MUAstabilityTheta (stab-θ)** |  |  |
| 7 | MUAstabilityAlpha |  |  |
| 8 | MUApower(raw)Delta | MUApowerDelta |  |
| 9 | MUApower(raw)Theta | MUApower(raw)Alpha | MUApower(raw)Beta |
|  | MUApower(raw)Gamma | MUApower(raw)HGamma |  |
| 10 | MUAstabilityBeta |  |  |
| 11 | MUAstabilityGamma |  |  |
| **12** | **MUAstabilityHGamma (stab-$\text{h}\gamma$)** |  |  |
| 13 | syncMUA | syncMUADelta | syncMUATheta |
| 14 | syncMUAAlpha |  |  |
| **15** | **MUApowerTheta (pow-θ)** | **MUApowerAlpha (pow-α)** |  |
| 16 | syncMUABeta |  |  |
| 17 | MUApowerBeta | MUApowerGamma | MUApowerHGamma |
| **18** | **syncMUAGamma (sync-γ)** |  |  |
| 19 | syncMUAHGamma |  |  |
| 20 | FringRatesstability |  |  |
| 21 | MUAstability |  |  |

Definitions of all features are detailed in the 'Signal processing and feature extraction' section of the Methods. Clusters No. 2, No. 6, No. 12, No. 15, and No. 18, which include key CT features significant for analysis—MUAstabilityTheta (stab-θ ), MUApowerTheta (pow-θ ), MUApowerAlpha (pow-α ), Firing Rates, syncMUAGamma (sync-γ ), and MUAstabilityHGamma (stab- $\text{h}\gamma$ )—are highlighted in bold.

**TableS3.** Feature weights and Stability-Importance index.

| Feature | aveWlow | aveWtop | Stable-Importance index (%) |
| --- | --- | --- | --- |
| **stab-θ** | **1.00** | **0.99** | **100** |
| **sync-γ** | **0.71** | **0.65** | **90** |
| **pow-θ** | **0.52** | **0.60** | **40** |
| pow-α | 0.65 | 0.36 | 10 |
| FiringRates | 0.62 | 0.35 | 10 |
| **stab-hγ** | **0.35** | **0.48** | **30** |

AveWlow: average feature weights within lowAcc (low accuracy) sets; aveWtop: average feature weights within topAcc (high accuracy) sets; stab-θ : MUA stability in the theta band; sync-γ : syncMUA in the gamma band; pow-θ : MUA power in the theta band; pow-α : MUA power in the alpha band; stab-hγ : MUA stability in the high gamma band; Those features that are included in the CT metric are highlighted in bold.

**TableS4.** Ablation study.

| Features weights |  |  |  |  |
| --- | --- | --- | --- | --- |
| stab-θ | 22.19 | 23 | 22.92 | 23 |
| pow-θ | 22.21 | 14.2 | 14 | |
| sync-γ | 15.58 | 13.44 | | |
| stab-hγ | 11.05 | | | |
| Performance |  |  |  |  |
| accuracy | 1 | 0.91 | 0.87 | 0.83 |
| F1 Score | 1 | 0.89 | 0.82 | 0.75 |
| AUC | 1 | 0.99 | 0.91 | 0.88 |
| p value | 0 | 0 | 0 | 0.01 |
| randAcc | 0.49 | 0.49 | 0.49 | 0.5 |

Stab- $\theta$ : MUA stability in the theta band; pow-θ : MUA power in the theta band; sync-γ : syncMUA in the gamma band; stab-hγ : MUA stability in the high gamma band; AUC: area under the curve value of the receiver operating characteristic; p value: the p value of Permutation Test; randAcc: baseline random accuracy.

**TableS5.** Network connectivity parameters.

| Connection | Type | $g_{E/I}$<br>(mS) | Connectivity | $\tau_s$ | $t_{\text{delay}}$ (ms) |
| --- | --- | --- | --- | --- | --- |
| inputs to TRN | Exc. | 0.01 | 1-to-1 | 10 | 7 |
| CT to TRN | Exc. | 0.01 | 1-to-many (Prob=0.15) | 20 | 3 |
| SR to TRN | Exc. | 0.02 | 1-to-1 | 20 | 3 |
| inputs to CT | Exc. | 0.0078 | 1-to-1 | 7 | 0 |
| TRN to CT | Inh. | 0.03 | 1-to-many (Prob=0.15) | 30 | 3 |
| inputs to SR | Exc. | 0.0078 | 1-to-1 | 7 | 0 |
| TRN to SR | Inh. | 0.02 | 1-to-1 | 30 | 3 |

TRN: Thalamic reticular nucleus; CT: Central Thalamus; SR: Specific relay nuclei; Exc.: Excitatory connections; Inh.: Inhibitory connections; $g_{E/I}$ : increase in synaptic conductance triggered when a presynaptic neuron fires; $\tau_s$ : a time constant that modulates synaptic conductance decay. $t_{\text{delay}}$ : delay time following presynaptic firing.

**TableS6.**
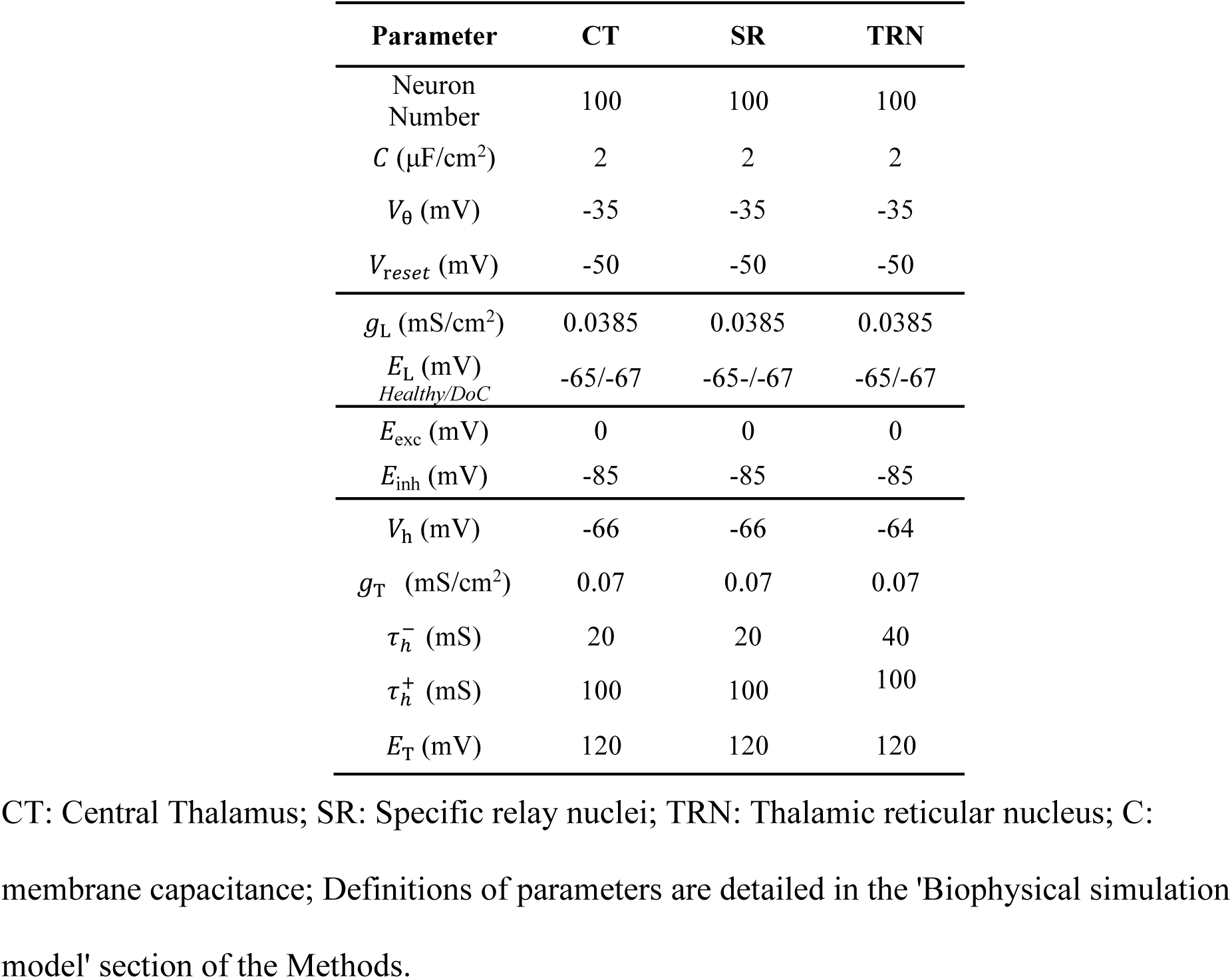
Thalamic neuronal parameters.

